# Advancements in monitoring: a comparison of traditional and application-based tools for measuring outdoor recreation

**DOI:** 10.1101/2024.02.09.579662

**Authors:** Talia Vilalta Capdevila, Brynn A. McLellan, Annie Loosen, Anne Forshner, Karine Pigeon, Aerin L. Jacob, Pamela Wright, Libby Ehlers

**Author notes:** Corresponding Author: Brynn A. McLellan^1^ Yellowstone to Yukon Conservation Initiative, 200-1350 Railway Ave, Canmore, Alberta, T1W 1P6.

## Abstract

Outdoor recreation has experienced a boom in recent years. While outdoor recreation provides wide-ranging benefits to human well-being and is an important feature of many protected and non-protected areas, there are growing concerns about the sustainability of recreation with the increased pressures placed on ecological systems and visitor experiences. These concerns emphasize the need for managers to access accurate and timely recreation data at scales that match the growing recreation footprint. Here, we compare spatial and temporal patterns of winter and summer recreation use using traditional and application-based tools across the Columbia and Canadian Rocky Mountains of western Canada. We demonstrate how recreation use can be estimated using traditional and application-based tools, although their accuracy and utility varies across space, season and activity type. Cameras and counters captured similar broad-scale patterns in count estimates of pedestrians and all recreation activities. Application-based data provided detailed spatiotemporal information on recreation use, but datasets were biased towards specific recreation types and did not represent the full recreation population. For instance, Strava Metro data was more suited for capturing broad-scale spatial patterns in biking than pedestrian recreation. Traditional tools including aerial surveys and participatory mapping captured coarser information on the intensity and extent of recreation, with the former tool capturing areas with low recreation intensity and the latter tool suited for capturing recreation information across large spatial and temporal scales. Application-based data should be supplemented with data from traditional tools including cameras or trail counters to identify biases in data and fill in data gaps. We provide a comparison of each tool for measuring recreation use, highlight each tools’ strengths and limitations, and suggest how to use these tools to address real-world monitoring and management scenarios. Our research contributes towards a better understanding of what tools are available to measure recreation and can help direct managers in selecting which tool, or combinations of tools, to use that can expand the rigor and scope of recreation research. This information can support decision-making and lead to the protection of ecological systems while allowing for high-quality recreation experiences.

**GRAPHICAL ABSTRACT:** 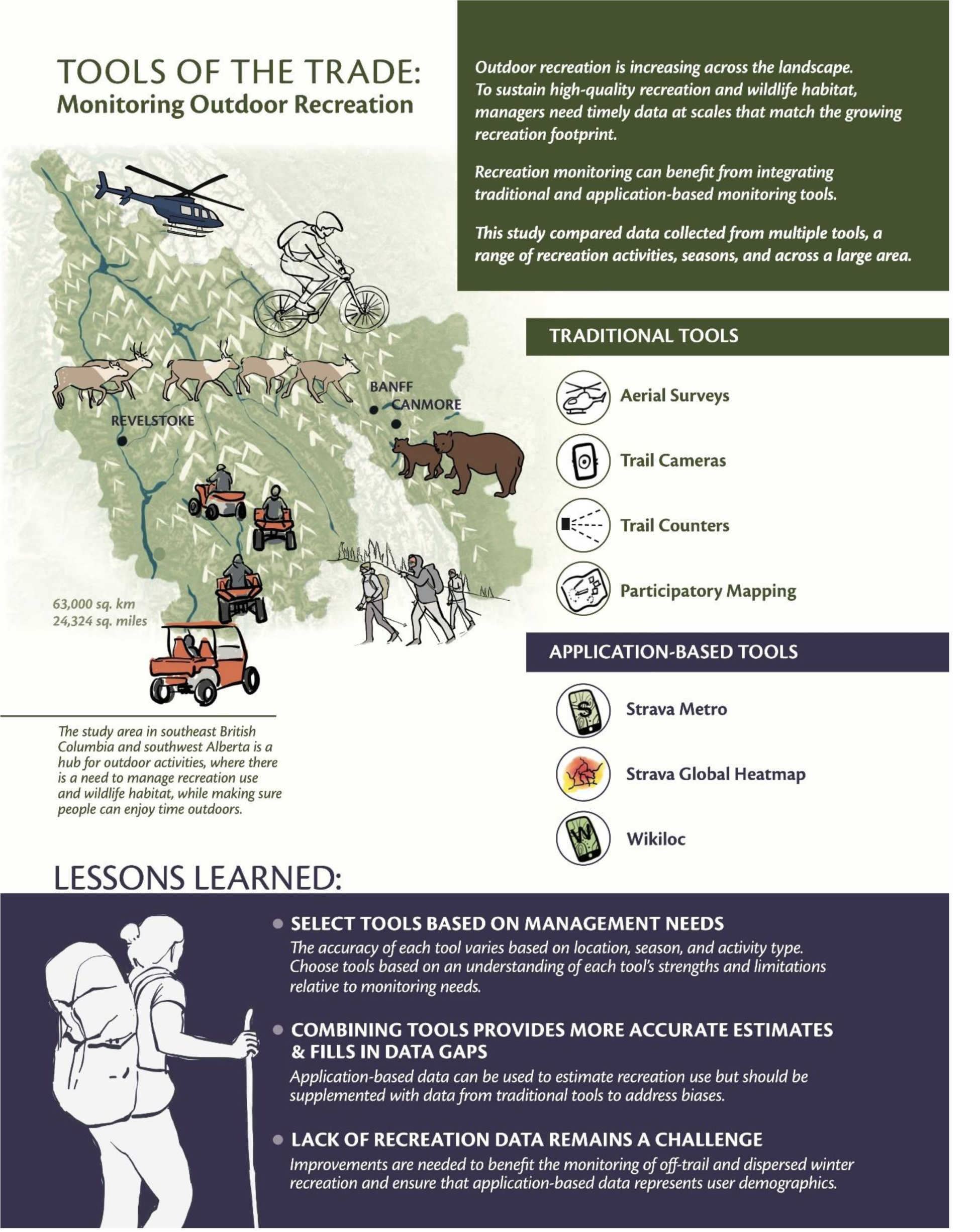

## 1. INTRODUCTION

Interest in outdoor recreation and visitation to parks, protected and non-protected areas is increasing globally, with terrestrial-protected areas receiving approximately eight billion visits annually (Balmford et al., 2015; Outdoor Industry Association, 2017). Outdoor recreation (hereafter, recreation) is an important element of many protected and non-protected areas and provides benefits to human well-being (Hartig et al., 2014; Lackey et al., 2021). However, there are growing concerns about the sustainability of recreation with the pressures that increasing recreation places on ecological systems (Larson et al., 2016; Schulze et al., 2018; Granados et al., 2023) and impacts to recreation experiences. Wildlife responses to recreation vary widely and are often context and species-specific (Larson et al., 2016; Granados et al., 2023). Recreation has been identified as the most common threat to species at risk in Canada (Rosenthal et al., 2022) and has been linked to reduced species richness, diversity and abundance (Sato, Wood & Lindenmayer, 2013; Larson et al., 2019), shifts in wildlife habitat use (Naidoo & Burton, 2020; Granados et al., 2023; Gump & Thornton, 2023), and loss and degradation of wildlife habitat (Heinemeyer et al., 2019). Simultaneously, the rise in recreation has led to crowding and conflicts between different user groups, reducing the quality of recreation experiences for many (Manning & Valliere, 2001; Jackson, Haider & Elliot, 2003; Wolf, Brown & Wohlfart, 2018; Ferguson et al., 2023).

As the demand for recreation grows, managers are increasingly faced with making decisions to protect plant and wildlife communities while supporting high-quality recreation experiences (Loosen et al., 2023). These decisions often necessitate understanding spatial and temporal patterns in recreation (Loomis, 2000) and the impact of recreation on ecological systems, including wildlife use patterns. Yet, an understanding of where, when, how intense, and what recreation activity types occur across a landscape remains difficult to attain. The lack of accessible, current, and spatiotemporal recreation data is consistently highlighted as a critical knowledge gap impeding managers’ decision-making and strategic planning (Loomis, 2000; Columbia Mountains Institute of Applied Ecology, 2023, Loosen et al., 2023). This hinders progress towards sustainable recreation (Hadwen, Hill & Pickering, 2007), especially for unpredictable and off-trail recreation activities in remote areas (Braunisch, Patthey & Arlettaz, 2011).

Numerous tools for measuring and monitoring recreation exist which address this data gap. In this study, we focus on two tool categories: traditional and application-based tools. We define traditional tools as any tool used specifically to monitor and count recreationists, and application-based tools as any tool that provides data from an activity sharing platform where geographic information is voluntarily shared by the app user (Goodchild, 2007). Generally, the primary purpose of application-based tools are centered around the user’s performance, fitness, or enjoyment rather than recreational research or monitoring. To achieve research, monitoring and management goals it is important to understand the advantages and limitations of each tool and their associated data (Cessford & Muhar, 2003; Norman & Pickering, 2017; McCahon, Brinkman & Klimstra, 2023). Understanding the differences between traditional and application-based tools has been recognized as a research priority (Norman & Pickering, 2017). While several studies have incorporated multiple tools to measure recreation (Norman & Pickering, 2017; Fisher et al., 2018; Wood et al., 2020), comparison of tools across large, continuous landscapes (i.e., inside and outside protected areas) and for many recreation activities remains relatively unexplored.

Traditional monitoring tools span a range of methods and include access permits, voluntary self-reporting or registration (Fredman et al., 2009), aerial surveys (Heinemeyer et al., 2019), on-site counts and surveys of people on trails or cars in parking lots (Fisher et al., 2018), participatory mapping (PM; Brown & Weber, 2011; Brown & Kyttä, 2014), and trail cameras and counters (Pettebone, Newman & Lawson, 2010; Miller, Leung & Kays, 2017; Naidoo & Burton, 2020). Many of these tools provide detailed and contextual information on the type of recreation activity and number, behavior or characteristics of recreationists. However, compliance of self-reporting permits and registration can vary due to a lack of enforcement and lead to underestimates of recreation use (Fredman et al., 2009). Moreover, on-site survey methods require prior information on where people are recreating, making it challenging to capture new areas (e.g., where people use trails or areas not managed for recreation; Vilalta Capdevila et al. 2022; Lawson 2021). Alternatively, managers and researchers can use PM to identify areas with recreation. PM engages participants to identify recreation areas or the intensity of recreation activities on a map (Brown & Weber, 2011; Rösner et al., 2014; Wolf et al., 2015; Wolf, Brown & Wohlfart, 2018; Komossa, Wartmann & Verburg, 2021). While PM can provide insight into the presence of and spatial distributions of recreation and recreationists’ behavior, cognitive bias can limit the accuracy and inference to specific times and locations (Brown, 2011). Other tools such as automatic infrared trail counters (hereafter, counters) and trail cameras (hereafter cameras) allow for the passive collection of georeferenced and time-stamped data (Lawson, 2021). Installing and maintaining large camera and counter networks across large landscapes can be logistically challenging and require substantial resources. For cameras, individual privacy must be protected if cameras are used for monitoring human recreation. Yet, large-scale studies are often needed to capture expanding recreation footprints and to characterize spatial and temporal recreation patterns at a scale appropriate for decision-makers (e.g., beyond a single protected park or trail network). As a result, traditional tools are often inadequate for collecting large-scale recreation data.

In contrast to traditional tools, application-based tools capitalize on large, crowdsourced datasets. These tools take advantage of the fitness and recreation data generated by the proliferation of mobile devices and global positioning systems (GPS), helping to overcome the challenges of many traditional tools (Wilkins, Wood & Smith, 2021). Application-based tools such as Strava, AllTrails, and Wikiloc are recreation-specific or fitness apps that collect geographical information when recreationists upload their activities (Norman & Pickering, 2017; Lee & Sener, 2020; Goodbody et al., 2021; Ferguson et al., 2023; Venter et al., 2023). Other social media apps such as Instagram, Flickr, and Twitter also provide detailed information of where, when and how people recreate (Heikinheimo et al., 2017; Fisher et al., 2018; Ghermandi & Sinclair, 2019).

Application-based tools link an individual’s activity track, photograph, or point location from these apps to a specific time and geographic coordinate. While application-based tools can generate high volumes of data across large areas and over extended periods (Fisher et al., 2018), analyses of these complex datasets often require technical know-how, extensive computational processing and storage (Lee & Sener, 2020; Wood et al., 2020) and consideration of privacy and ethical use. Further, application-based tools can also be spatially and temporally biased in the type of users, geography, and data available (Norman & Pickering, 2017; Venter et al., 2023).

Despite the promise of application-based tools for measuring recreation, these types of data require careful consideration of biases and assumptions that are beginning to be explored (Venter et al., 2023).

If multiple tools for monitoring recreation are to be adopted, there is a need to compare and understand the different tools and data sources to provide clear recommendations for guiding evidence-based management and planning for recreation, land use and wildlife. To address this need, we compare data from traditional and application-based tools to measure spatial and temporal patterns of summer (May 1 – Oct 31) and winter (Nov 1 – Apr 30) recreation use. We use data from four traditional tools (counters, cameras, aerial surveys, PM) and three application- based tools (Strava Metro, Strava Global Heatmap, Wikiloc) across western Canada. This area is a hub for motorized and non-motorized recreation and there are growing concerns about managing conflicts between recreation use (Holterman, Wright & Jacob, 2023) and maintaining high-quality wildlife habitat. We explore the relationships between recreation data from tools over time, space, and recreation activity types (Table 1). First, we compare recreation data across the study area to examine broad use patterns across tools. Second, we use case studies to assess smaller areas and trail network-specific use patterns. Collectively, this work provides a unique opportunity to compare data collected across multiple tools, a range of recreation use intensities and activity types, and a large spatial extent that captures a range of human development and ecological conditions.

**Table 1.** Research questions comparing data from traditional and application-based tools.

|  | Research question | Tools compared | Recreation type | Season<sup>1</sup> |
| --- | --- | --- | --- | --- |
| **Study area** |  |  |  |  |
| **1A & 1B** | How do monthly counts of recreation use correlate across space (1A) and time (1B?). | Strava Metro<sup>2</sup>, cameras and counters | all activities, biking, pedestrian<sup>5</sup> | year-round |
| **2** | How do annual counts of recreation use correlate across space? | Strava Heatmap<sup>3</sup>, Strava Metro<sup>2</sup>, cameras, counters | all activities | year-round |
| **Case study** |  |  |  |  |
| **1** | How do annual counts of recreation use correlate across space? | Wikiloc, PM<sup>4</sup> | motorized | summer |
| **2** | How does the spatial distribution and intensity of winter motorized recreation use compare across space? | aerial surveys, Wikiloc, PM<sup>4</sup> | motorized | winter |
| **3** | How do monthly counts of biking recreation use correlate across space? | Strava Metro<sup>2</sup>, cameras, counters | biking | year-round |

<sup>1</sup> Season dates: summer (May 1 – Oct 31) and winter (Nov 1 – Apr 30).
<sup>2</sup> Strava Metro contains counts of recreation users by activity type (biking, pedestrian) and is only available to Strava Metro partners.
<sup>3</sup> Strava Heatmap contains aggregated annual indices of recreation (run, hike, walk, ride, water, winter) and is publicly available.
<sup>4</sup> Participatory mapping (PM).
<sup>5</sup> Pedestrian includes foot activities such as hiking, walking, running.

## 2. METHODS

### 2.1 Study Area

The study area encompasses 63,000 km^2^ of rugged mountainous terrain in western Alberta and eastern British Columbia, Canada (Fig. 1), and contains 20,260 km^2^ (32%) of federal and provincial protected areas, including provincial and federal parks, wilderness areas, and heritage rangelands (Loosen et al., 2023). The region includes open public and private lands, grazing leases, motorized (e.g., public land use zones in Alberta) and non-motorized recreation areas (e.g., National Parks). Commercial snowmobiling and heli-assisted activities occur in some areas not serviced by maintained roads. The study area includes the population centers of Canmore, Banff, Golden, Revelstoke, Nakusp, and Invermere (Fig. 1).

**Figure 1.**
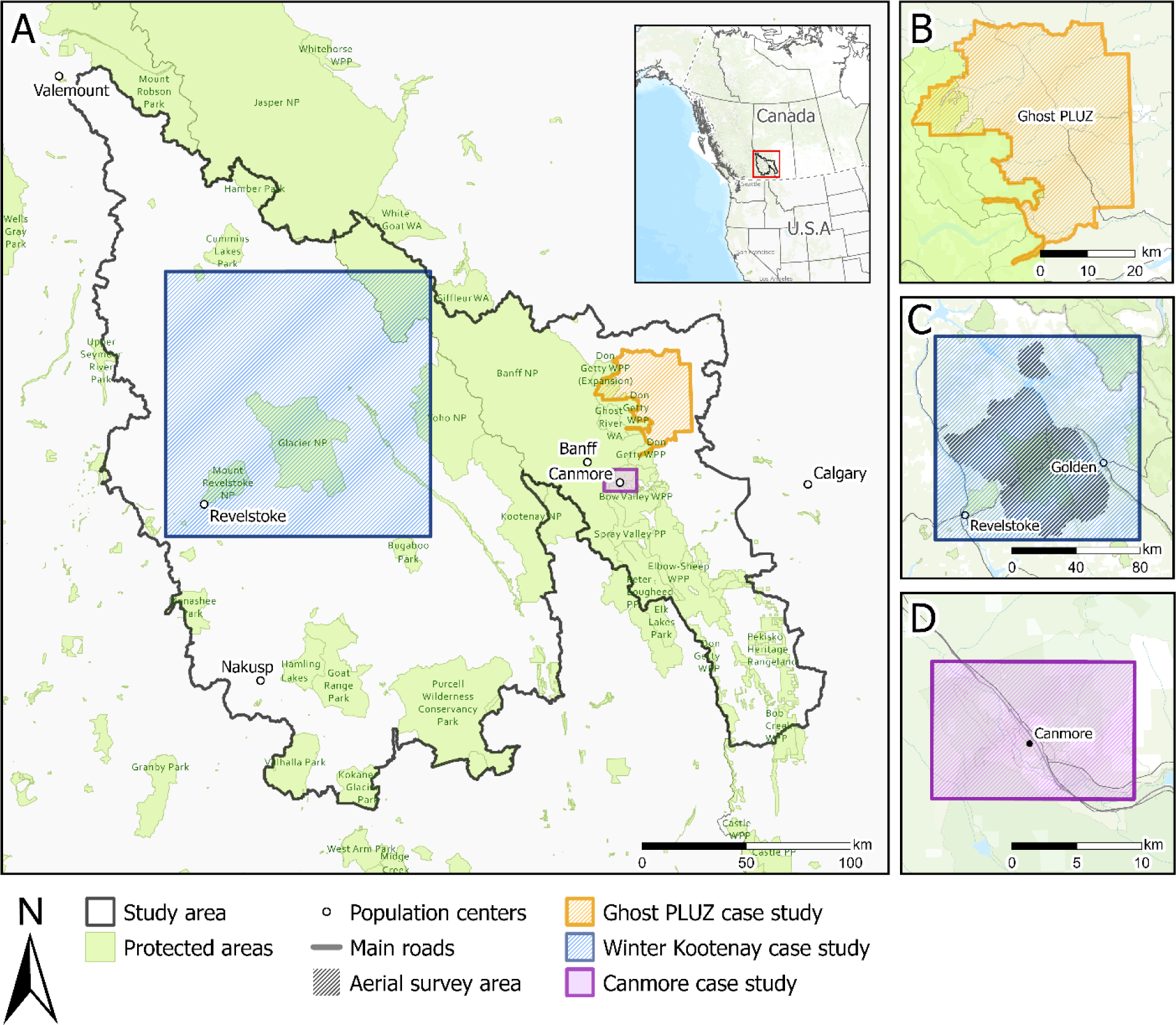
**Study area (panel A) and case study areas (panels B, C, D)**. Case studies include the Ghost Public Land Use Zone (PLUZ, panel A) that has a high density of motorized trails in a non-protected area, the Kootenay mountains (panel C) that has winter motorized recreation, and biking trails surrounding the town of Canmore in the Bow Valley (Panel D).

The study area is within the traditional territories of the Okanagan/Syilx, Sinixt, Ktunaxa, Secwépemc, Niitsitapi (Blackfoot) Nations of Siksika, Kainai, Piikani, and Aamskapi Pikuni; the Îyârhe (Stoney) Nakoda Nations of Goodstoney, Bearspaw, and Chiniki; Tsuut’ina First Nation; Mountain Cree. It also includes lands within Treaties 6, 7, and 8 and districts 1, 3 and 4 of the Otipemiskiwak Métis Government of Alberta. The study is part of the greater Yellowstone to Yukon region that includes key linkages for ecological connectivity, core protected areas and human modified landscapes (Holterman et al., 2023). This region includes alpine, sub-alpine and montane subregions and habitat for mammals sensitive to human presence such as grizzly bears (Ladle et al., 2018), wolverines (Heinemeyer et al., 2019; Barrueto et al., 2022) and caribou (Seip, Johnson & Watts, 2007; Wilson & Wilmshurst, 2019).

Resource developments are prominent across the study area and include forestry, mining and oil and gas. Forestry and mining operations contribute to polygonal footprints and resource roads (Forest Practices Board, 2021). The study area includes approximately 53,436 km of trails, gravel resources roads, and paved roads with an overall density of 0.85 km/km^2^ (Loosen et al., 2023). Roads and trails associated with resource and industrial developments provide access points for motorized and non-motorized recreation (Ministry of Forests, Lands, and Natural Resource Operations, 2013; Government of Alberta, 2018; Forest Practices Board, 2021).

### 2.2 Measuring Recreation

#### 2.2.1 Traditional Tools

##### Trail Counters

We acquired data from TRAFx Infrared Trail or Vehicle Counters (TRAFx Research Ltd., Canmore, Alberta, Canada) from partners (provincial and federal government, researchers, and not-for-profit organizations). Counters were positioned along trails and roads to count the number of people or vehicles that pass through the device’s infrared scope. Details on counter models and set up are provided in supplemental information.

In ArcMap (ArcMap Version 10.8.2, ESRI, Redlands California) we retained data from trails only (Vilalta Capdevila et al., 2022) and filtered by activity type. A portion of these devices count bicycles (vehicle counters with bike mode activated) and we considered any data from bike counters to represent bike counts only, while data from the other counters were assumed to represent pedestrian counts. TRAFx trail counters do not differentiate between recreation activity types, and we could not validate if any other activities were counted. To represent recreation intensity, count data was aggregated from daily counts to monthly totals per year from 2017 to 2019 for pedestrian, biking, and all recreation activities.

##### Cameras

We obtained camera datasets from provincial and federal government and researcher partners. Remote trail cameras (hereafter, cameras) were deployed on trails and low-traffic roads to monitor wildlife but also to collected recreation data. Camera images were classified by the number of recreationists and recreation activity type. To represent recreation intensity, we aggregated daily counts of recreationists into monthly totals per year from 2017 to 2019 in three categories: pedestrian (any foot-based activity) and biking (included road, mountain, electric and fat bikes), and all activities (includes all motorized and non-motorized activities). Details on activity types, camera models, set up, and image classification are provided in supplemental information.

##### Aerial surveys

We conducted aerial surveys to quantify the intensity and spatial extent of winter recreation, that includes off and on-trail use, in three survey areas within the study area (Fig. 2; Fig. S3). We completed two rounds of surveys in a helicopter between February and April 2022, with each survey occurring at least a month apart. Following Heinemeyer et al. (2019), we organized the survey into 1.5 x 1.5 km (2.25 km2) grids. We flew transects, that were spaced 3 km apart, with two observers seated on either side of a helicopter. We recorded the type of recreation (snowmobiling and backcountry, heli and cat skiing) and the spatial footprint of recreation tracks within a 1.5 x 1.5 km observation window. We classified the footprint into five bins representing the percent cover of recreation tracks: <10, 10–25, 26–50, 51–75, >76%). Raw data were processed to convert observation waypoints into a polygon grid representing the observation window; see supplementary information for further details.

**Figure 2.**
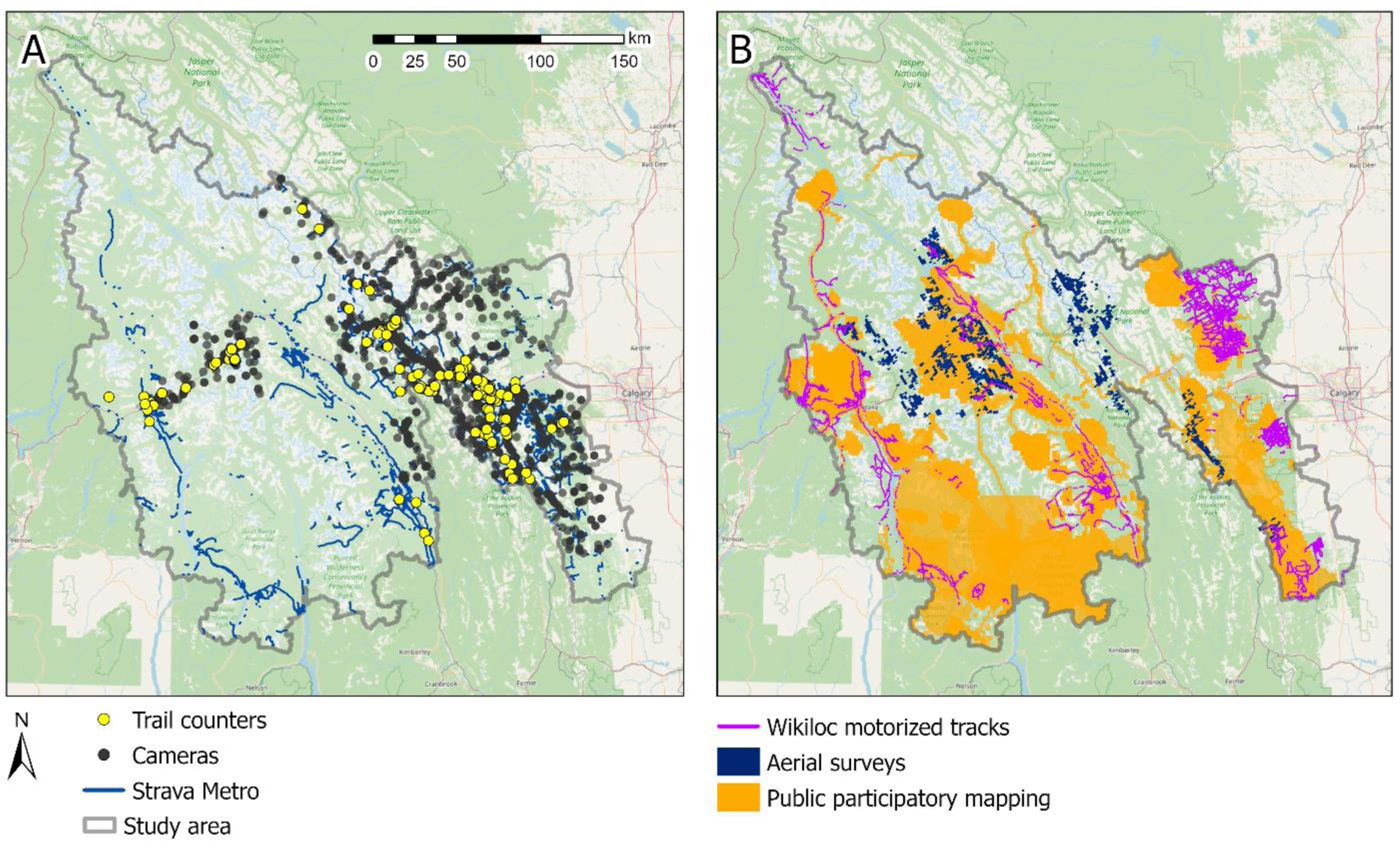
Spatial distribution of data from counters, cameras, Strava Metro (panel A) and public participatory mapping, aerial surveys, and Wikiloc (panel B) within the study area (gray boundary)**. Strava Heatmap not displayed (see Fig. S5).**

##### Participatory mapping

We conducted participatory mapping (PM) interviews with recreation experts (e.g., park rangers, recreation group, trail users, lodge and campground owners) across our study area (Fig. 2). Meetings were conducted from 2020 to 2021 over Zoom (Zoom Video Communications, Inc. 2020, San Jose, California United States). We shared a digital map on the screen and asked participants to identify areas with recreation and the types of activities and average number of people per day recreating in these areas from 2017 to 2019 (Fig. S2).

Participants marked locations on the shared map. See supplementary information for further details. We categorized the estimated daily number of people recreating in an area into three bins: 0–10; 11–50; > 51 recreationists per day. Recreation areas were digitized to a 1.5 x 1.5 km cell raster. Each cell was assigned one of the three bin values to represent recreation intensity. In areas where participants indicated differing numbers of recreationists in the same place, we used the highest estimate.

#### 2.2.2 Application-based tools

##### Strava Metro

Strava Metro (https://metro.strava.com) uses non-motorized biking and pedestrian (running, walking, hiking) GPS data collected from users’ phones, fitness watches or GPS devices. Strava Metro snaps GPS tracks to the closest OpenStreetMap (www.openstreetmap.org) street and trail segment. OpenStreetMap is a crowdsourced geographical database of the world. Strava Metro data are anonymized to maintain user privacy by aggregating count data for trail segments with at least three unique users per direction (up or down the activity segment), representing totals as multiples of five (Gamez, 2021, pers. comm.). Strava Metro provides data as hourly, daily, monthly, or yearly counts. We aggregated the number of users in each direction along a trail segment into monthly totals per year from 2017 to 2019 for biking, pedestrians and all activities. We acquired a Strava Metro partnership through our work with Alberta and British Columbia government ministries.

##### Strava Global Heatmap

Strava Global Heatmap (hereafter, Strava Heatmap; https://www.strava.com/heatmap) is a crowdsourced heatmap that uses GPS-based recreation tracks of public non-motorized recreation activities (run, hike, walk, ride, water, winter) recorded on phones, fitness watches, and GPS devices. Strava Heatmap is freely available, unlike Strava Metro, and has been used as an index of human recreation (Jäger, Schirpke & Tappeiner, 2020; Corradini et al., 2021; Carlson et al., 2022). The heatmap displays annual aggregated indices of public activities for each calendar year, with updates to the heatmap occurring monthly. The Strava Heatmap visualizes the accumulation of individual user tracks and uses a cumulative distribution function to normalize the intensity of “heat” that maximizes contrast. The map also has a bi-linear interpolation applied to avoid visual artifacts in areas of low use or large gradients of use (Robb, 2017). Roads and trails with little use are not displayed on the heatmap until several tracks are uploaded. We used the 2016 to 2017 heatmap and converted the raster to a greyscale gradient such that the color of each cell represented an index of recreation intensity, unlike Strava Metro that provides counts of app-users tracks. We converted the data using the ‘greyscale’ image analysis function in ArcGIS Pro (Version 3.1.0) to produce a grid of high and low use intensity areas (max = 255, min = 0).

##### Wikiloc

Wikiloc (https://www.wikiloc.com) uses GPS-based recreation tracks recorded on phones, fitness watches, and GPS devices that users can upload to a public website. Wikiloc data includes a range of motorized and non-motorized recreation activities. We downloaded motorized tracks (quad, snowmobile, off-road vehicle, motorbike) available within our study area (Fig. 2). We saved the GPS tracks, recreation type, time and date for each track and removed any unique personal identifiers of users to maintain data privacy.

### 2.3 Analyses

#### 2.3.1 Comparison of monthly counts across time and space

We correlated counts from cameras, counters, and Strava Metro across space and time (questions 1A and 1B; Table 1). Datasets were acquired from multiple research projects and therefore were not always aligned spatially (Fig. 2) or temporally (Table 2). To match tools, devices were paired when cameras and counter were within 200 m of each other along the same trail or were within 30 m of a Strava Metro trail segment and had data for the same day. To match cameras and counters, we used trails to create a distance raster between devices and selected distances within 200 m and data from the same day. We selected the distance threshold of 200 m so that each tool was likely to capture the same recreationist along a trail, while maximizing the number of matched locations. We used 30 m to match the trail segments to the camera or counter because Strava Metro data are already linked to OpenStreetMap trail segments and the buffered distance used to create the trail network dataset used in this work was 30 m (Vilalta Capdevila et al., 2022). We refer to matched pairs of tools as matched locations. We grouped analyses into all recreation activity types, pedestrian, and biking. We did not include comparisons of motorized activities due to data sparsity.

**Table 2.** Comparison of tools to monitor recreation use. . The table includes a description of how each tool quantifies recreation use, data type (resolution), minimum temporal grain, temporal and spatial extent, recreation activity type (motorized, non-motorized, all recreation activity types), season (summer, winter, year-round).

| **Tool** | **How recreation is quantified** | **Data type** | **Temporal grain** | **Temporal extent** | **Spatial extent<sup>1</sup>** | **Activity** | **Season** |
| --- | --- | --- | --- | --- | --- | --- | --- |
| **Counters** | number of recreationists per unit time | point | daily | 2017 - 2019 | portion | all activities | year round |
| **Cameras** | number of recreationists per unit time | point | daily | 2017 - 2019 | portion | all activities | year round |
| **Aerial surveys** | percent cover of recreation tracks | grid (1500 x 1500m) | daily | 2022<br>Feb - Apr | portion | all activities | winter |
| **PM<sup>2</sup>** | number of recreationists per area | grid (1500 x 1500m) | annual | 2017 – 2019 | portion | all activities | year round |
| **Strava Metro** | number of trail segments per unit of time | linear | daily | 2017 - 2019 | full study area | non-motorized | year round |
| **Strava Heatmap** | index of recreation intensity | grid (19 x 19m) | annual | 2016 - 2017 | full study area | non-motorized | year round |
| **Wikiloc** | number of trail segments per unit of time | linear | daily | 2013 - 2023 | portion | motorized | year round |

<sup>1</sup> See Fig. 2 for a map of the spatial distribution of the different tools.
<sup>2</sup> Participatory mapping (PM).

To compare counts between tools across space, we calculated Pearson’s correlation values for the monthly counts (sum of counts within a month) for all matched locations across three years (2017, 2018, 2019) for all activities, pedestrian, and biking. To compare counts between tools over time, we calculated Pearson’s correlation for the monthly counts for each of the three years for all activities, pedestrian, and biking. We considered correlation values greater than 0.75 as strong correlation. All analysis occurred in R Version 4.3.1 (R Core Team, 2021)

#### 2.3.2 Comparison of annual counts across space

We compared data from cameras, counters, and Strava Metro to aggregated indices of recreation use Strava Heatmap (question 2; Table 1). We visually compared annual Strava Heatmap index values to median annual counts from Strava Metro and cameras and counters that were deployed for a minimum of nine months (273 days) within a year. We chose nine months as our temporal unit for the cameras and counters to best match Strava Heatmaps temporal scale (i.e., the heatmap represents the annual aggregated indices of public activities) while at the same time including a range of camera and counter devices across the study area because most devices, particularly cameras, did not collect data continuously across a year. For Strava Metro, we compared the heatmap intensity values to the mid-point of the nearest Strava Metro segment. For cameras and counters, we generated a 50 m buffer around each device to match the maximum width of trails and used the highest raster value from Strava Heatmap within the buffer as a relative index of recreation intensity. This approach accounted for the heatmap displaying high use trails with a wider footprint relative to low use trails.

## 3. RESULTS

We recorded 2,030,653 total recreation counts from cameras, 79,775,734 counts from counters, and 28,211,105 counts from 32,374 Strava trail segments. For all recreation activities we compared 58 spatially matched locations of cameras and counters, 156 spatially matched locations of cameras and Strava Metro and 189 spatially matched locations of counters and Strava Metro. See Table S1 for spatially matched locations for biking and pedestrian activities. Some counters displayed extreme monthly counts for pedestrian and biking activities and all tools displayed generally higher counts in the spring and summer (Fig. S6). We collected 1673 unique Wikiloc motorized tracks from 2013 – 2023 within the study area. PM data was collected from 36 participants (see supplemental information for further details).

### 3.1.1 Comparison of monthly counts across space and time

#### Spatial correlation

There were high correlation values across spatially matched counters, cameras and Strava Metro (minimum mean correlation of 0.72 across all comparisons; Fig. 3). Mean correlation was highest for all activities, compared to pedestrian or biking (Fig. 3). Mean correlation values were the highest for the cameras and counter comparisons for all activities (r = 0.92; Fig. 3) and pedestrians (r = 0.91; Fig. 3). Correlation values of all activities, pedestrians and bikers varied across the study area, with some clusters of spatially paired counters, cameras and Strava Metro locations with low correlation (r < 0.5) surrounding the developments of Canmore, Banff and Lake Louise (Fig. S7).

**Figure 3.**
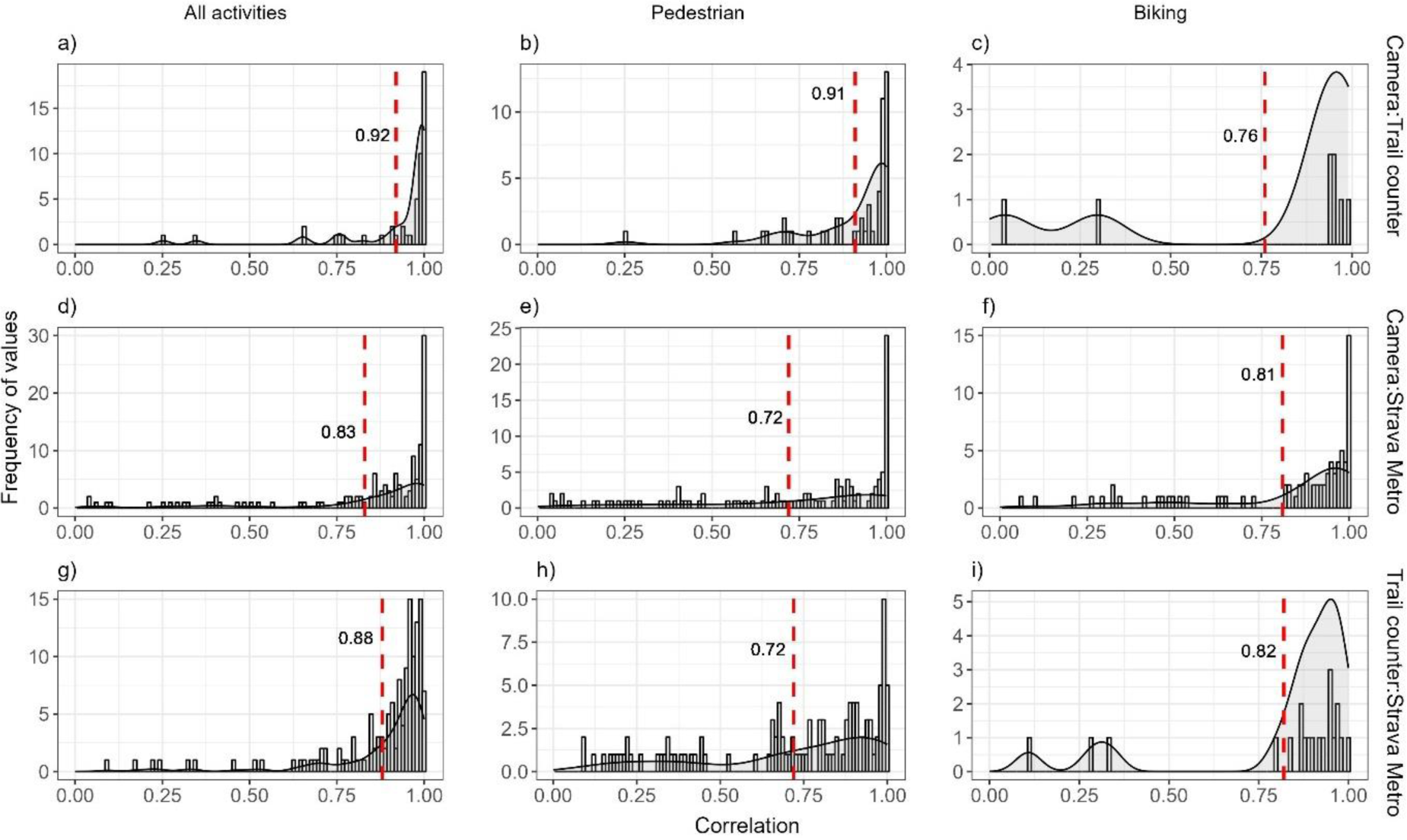
Distribution of Pearson’s correlation values of monthly counts of all recreation activities (A), pedestrian (B), and biking (C) for pairwise combinations of spatially matched cameras, counters and Strava Metro, 2017 – 2019. Each row represents different data comparisons and the dashed red line represents the mean Pearson’s correlation value.

#### Temporal comparison

Across all tools Pearson’s correlation values varied by month and year (Fig. 4). Low correlation values occurred generally during May to August, with this pattern most pronounced for comparisons between Strava Metro and cameras or counters for all activities (Fig. 4). Correlations for camera and counters generally remained consistently high year-round for all activities and pedestrians (Fig. 4). Biking displayed more variable correlations across months and years compared to all activities and pedestrians (Fig. 4).

**Figure 4.**
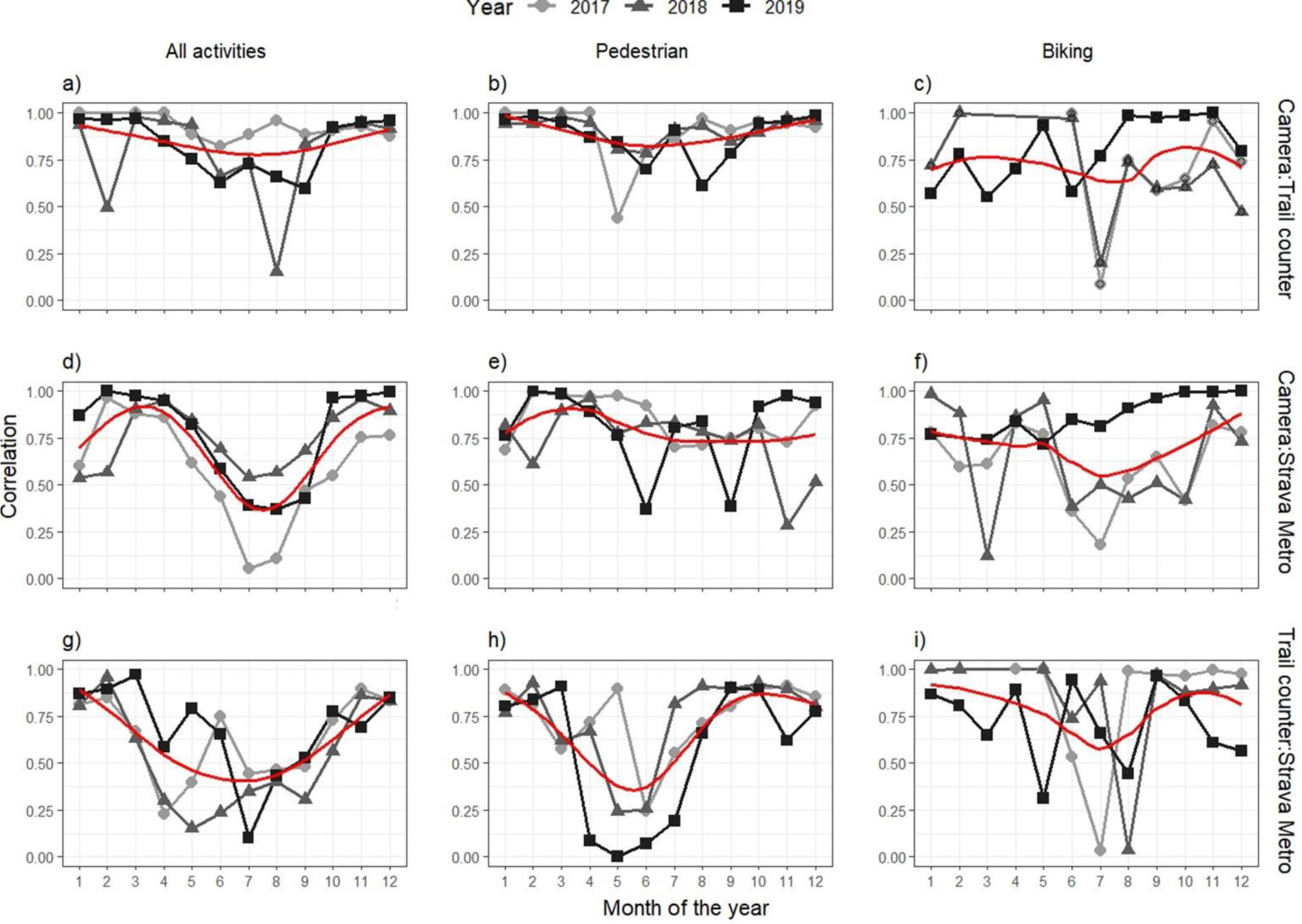
Pearson’s correlation values for monthly counts for all recreation activities (A), pedestrian (B), and biking (C) for pairwise combinations of spatially matched cameras, counters and Strava Metro, 2017 – 2019. The red horizontal line represents a non-linear trendline (generalized additive model).

### 3.1.2 Comparison of annual counts across space

High Strava Heatmap values coincided with both high and low annual median counts of cameras, counters or Strava Metro (Fig .5). There were many locations where high heatmap values coincided with low median counts of the three tools, particularly for counters and Strava Metro (Fig. 5). There were also locations where Strava Heatmap indicated no recreation yet cameras and Strava Metro captured recreation (Fig. 5).

**Figure 5.**
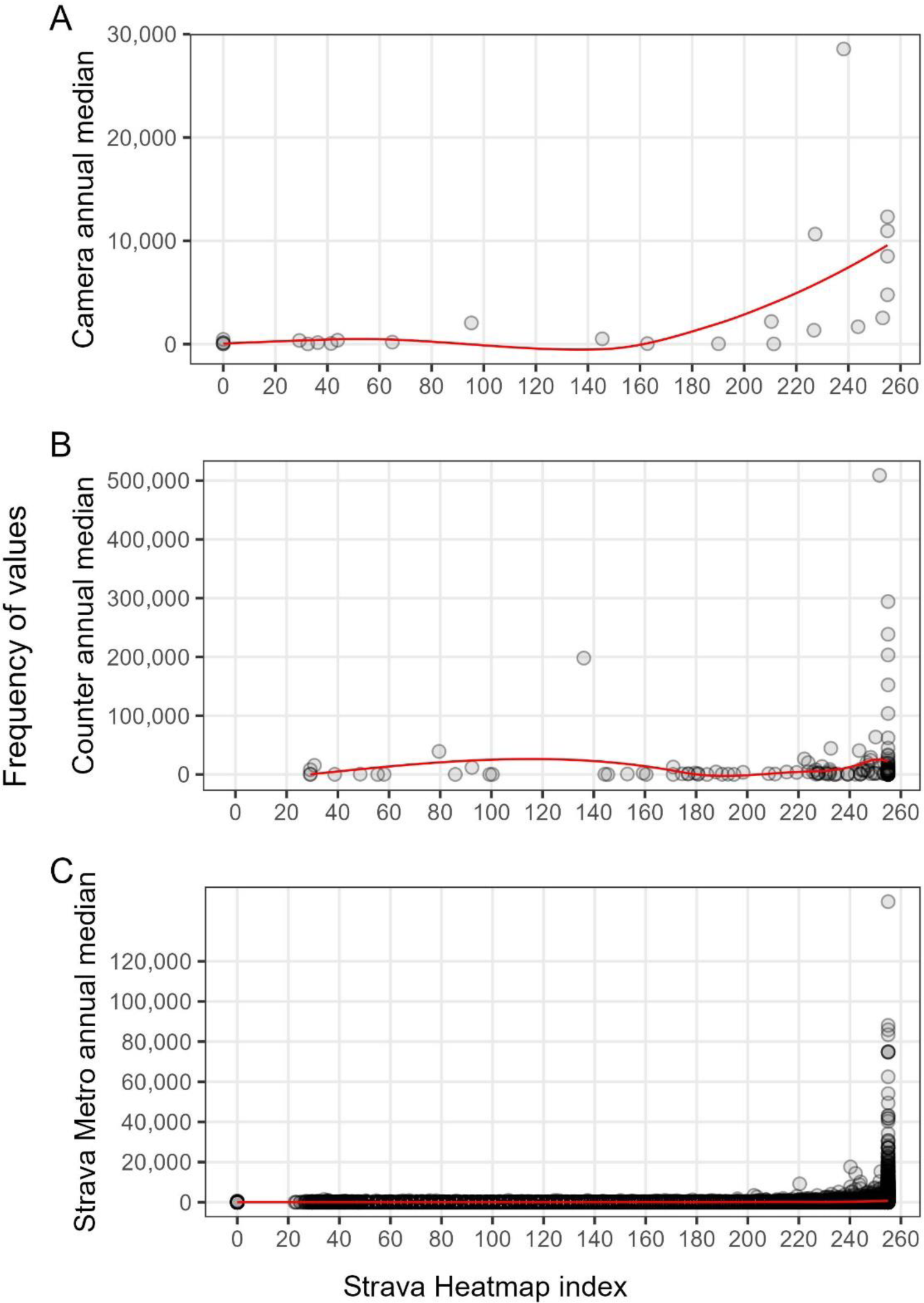
Strava Heatmap index of recreation use compared to annual median recreation counts from cameras (A), counters (B), and Strava Metro (C). Data for cameras, counters and Strava Metro are from 2017 to 2019 for all spatially matched locations in the study area, data from Strava Heatmap is from 2016 to 2017. Trendline represented in red.

## 4. CASE STUDIES

### 4.2 The Ghost Public Land Use Zone

Assessing the distribution and intensities of off-trail recreation remains a long-standing challenge in land-use and recreation management and planning. While traditional tools such as counters or cameras can provide spatiotemporal information of recreation on trails, it can be difficult for these devices to capture recreation across large areas and trail networks with many entry points or where use is dispersed (e.g. backcountry skiing in alpine areas). Participatory mapping (Brown & Weber, 2011; Wolf et al., 2015; Komossa, Wartmann & Verburg, 2021) and application-based data (Norman & Pickering, 2017; Ghermandi & Sinclair, 2019) offer alternative approaches for collecting recreation data, but it remains unclear how well the data reflects use of the total recreation population and how the data from different tools complement each other.

Here we compare the use of PM and application-based data from the recreation app Wikiloc for estimating summer motorized recreation use in the Ghost Public Land Use Zone (PLUZ), Alberta, Canada (Fig. 1; Fig. 6; Fig. 7). This area includes more than 1,500 km^2^ of public lands east of Don Getty Wildland Provincial Park and Banff National Park, Alberta (Government of Alberta, 2018) and is composed of a large network of designated and non-designated motorized trails (Yarmoloy & Stelfox, 2011; Weerstra, 2018; Fig. 1). The area is used for forestry, agriculture, oil and gas, and hosts a range of year-round recreation activities, including a designated trail network providing trails to OHVs specific to vehicle types and sizes. There are several designated provincial campgrounds and opportunities for un-serviced camping in designated camping nodes and random camping. To evaluate annual counts of summer motorized use between tools, we aggregated binned PM estimates of the number of summer motorized recreationists into 1.5 x 1.5 km grid cells. To match the annual time scale of the PM datasets, we partitioned the Wikiloc data from 2017 to 2019 to include motorized activities from May to September. We calculated the total length of tracks within each cell to produce a density (km of tracks/km^2^) grid in ArcMap.

**Figure 6.**
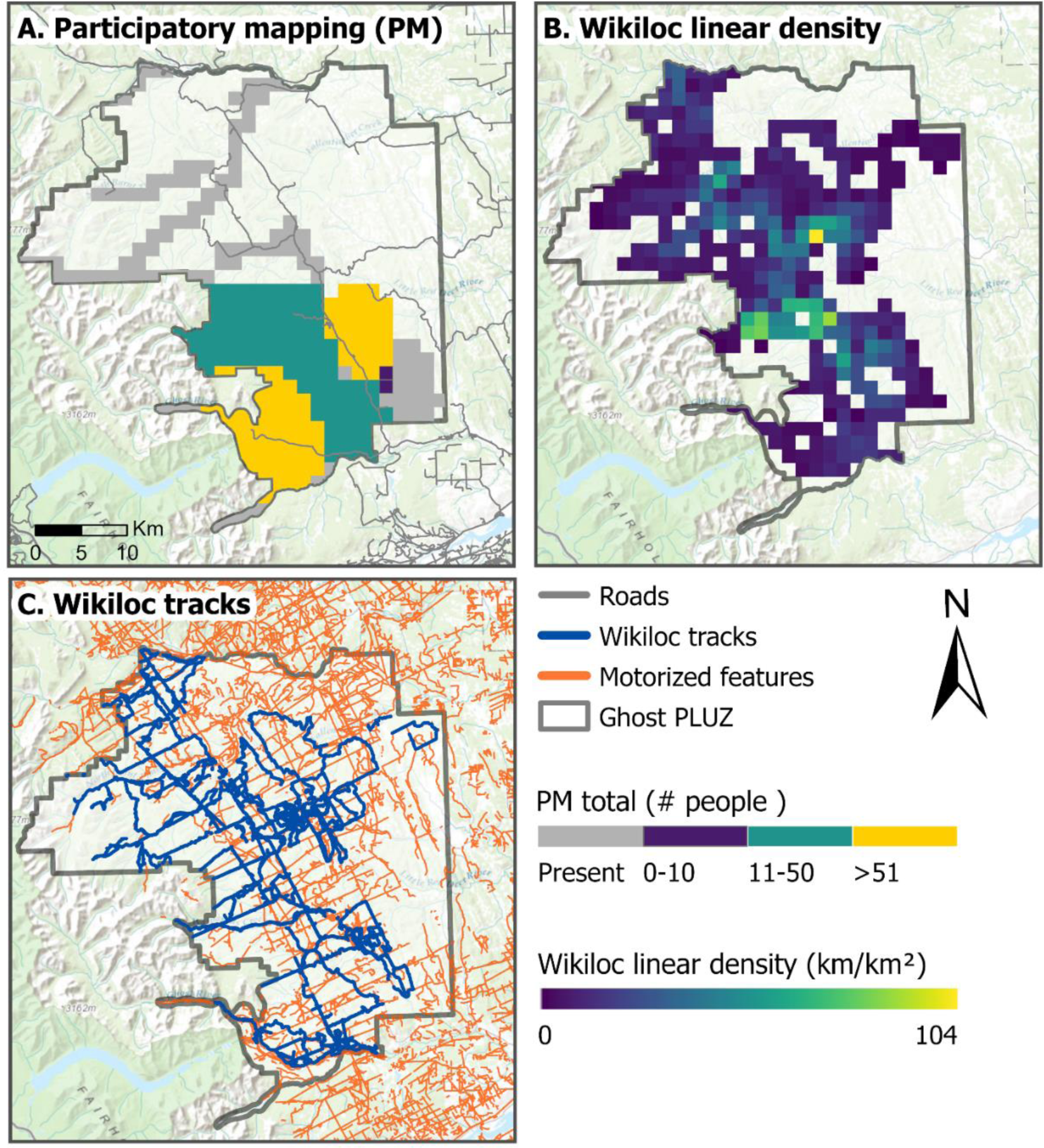
Summer motorized recreation in the Ghost Public Land Use Zone (PLUZ), Alberta from May to September, 2017 – 2019. Panel A represents estimates of the average number of people from participatory mapping (PM); gray shading represents areas where PM participants indicated that motorized recreation was present, but no count estimates were provided. Panel B displays the density of total motorized Wikiloc tracks (km/km^2^). Panel C displays Wikiloc tracks and motorized trails (Vilalta Capdevila et al., 2022).

**Figure 7.**
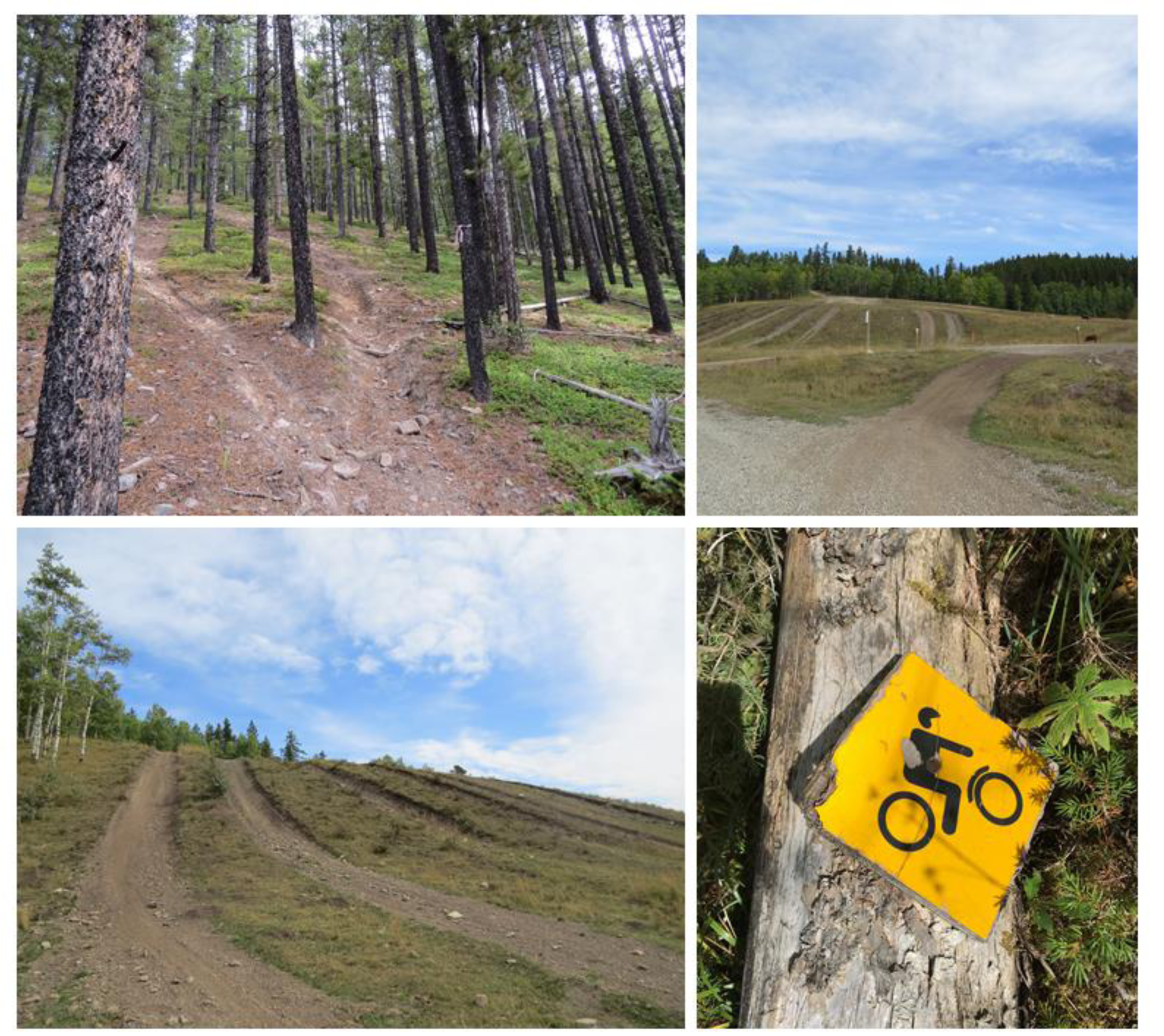
Recreation trails in the Ghost Public Land Use Zone, Alberta, Canada, September 2023. Photo credit: Talia Vilalta Capdevila, Brynn McLellan.

Both tools captured similar spatial extents of recreation, although patterns in recreation use intensity varied due to the resolution and user-group biases of the datasets (Fig. 6). Although Wikiloc provides spatiotemporal information of on and off-trail recreation use in remote areas, the data represent the relative recreation use among app users and therefore is unlikely to be representative of all recreationists (Norman & Pickering, 2017). Here we found that the Wikiloc data did not capture all motorized trails in the Ghost PLUZ (Fig. 6). Comparatively, PM provided information on the spatial extent and the approximate number of recreationists in some areas of the Ghost PLUZ. Estimates of use were not provided for all areas in the Ghost PLUZ, however, in these areas, some PM participants indicated that un-designated trails greatly outnumbered designated trails (Fig. 6). The combination of general and area-specific information generated from PM and Wikiloc datasets illustrate how combining these tools allow for a more robust assessment of recreation use and could be used to strategically identify recreation areas to monitor with traditional tools. This approach would be particularly useful in large areas with spatially dispersed or high-density trails systems with many entry point, such as the Ghost PLUZ, where employing and maintaining traditional tools can be time-intensive and costly (Wolf, Brown & Wohlfart, 2018).

### 4.3 Winter Motorized Recreation in the Kootenay Mountains

Backcountry winter recreation is a fast-growing recreation sector, with many rural mountain towns relying on winter recreation for their local economic activity (Outdoor Industry Association, 2017). The growing popularity of winter recreation combined with advancements in recreation equipment has allowed recreationists to travel further into remote and undisturbed areas, including sensitive wildlife habitats (Heinemeyer et al., 2019). Recreational activities can displace snow-dependent species from high quality habitats, such as wolverine (Heinemeyer et al., 2019), caribou (Seip, Johnson & Watts, 2007) and mountain goats (Richard & Côté, 2016).

Managing winter recreation to allow for the protection of wildlife habitat remains an ongoing challenge, one that is further intensified by the lack of data on dispersed, off-trail winter recreation. It can be difficult to capture dispersed winter-use using many traditional tools, such as cameras or counters, because snowpack enables widespread travel either on-or off-trails where fallen trees and shrubby vegetation are covered.

To highlight the challenges of monitoring off-trail winter recreation we compare the use of participatory mapping (PM), aerial flight surveys and application-based data from the recreation app Wikiloc for capturing winter motorized recreation near the population centers of Golden and Revelstoke in the Kootenay Mountains, British Columbia, Canada (Fig. 1; Fig. 8). These population centers are hubs for winter recreation and tourism and rely on recreation for their local economies (Morten & Der, 2019; Tourism Revelstoke 2019). We filtered the PM and aerial survey datasets to include snowmobiling and the Wikiloc dataset to include motorized activities from November to April, 2015 – 2023, and visually compared the spatial extent and use intensity of winter motorized recreation activities across the three tools.

**Figure 8.**
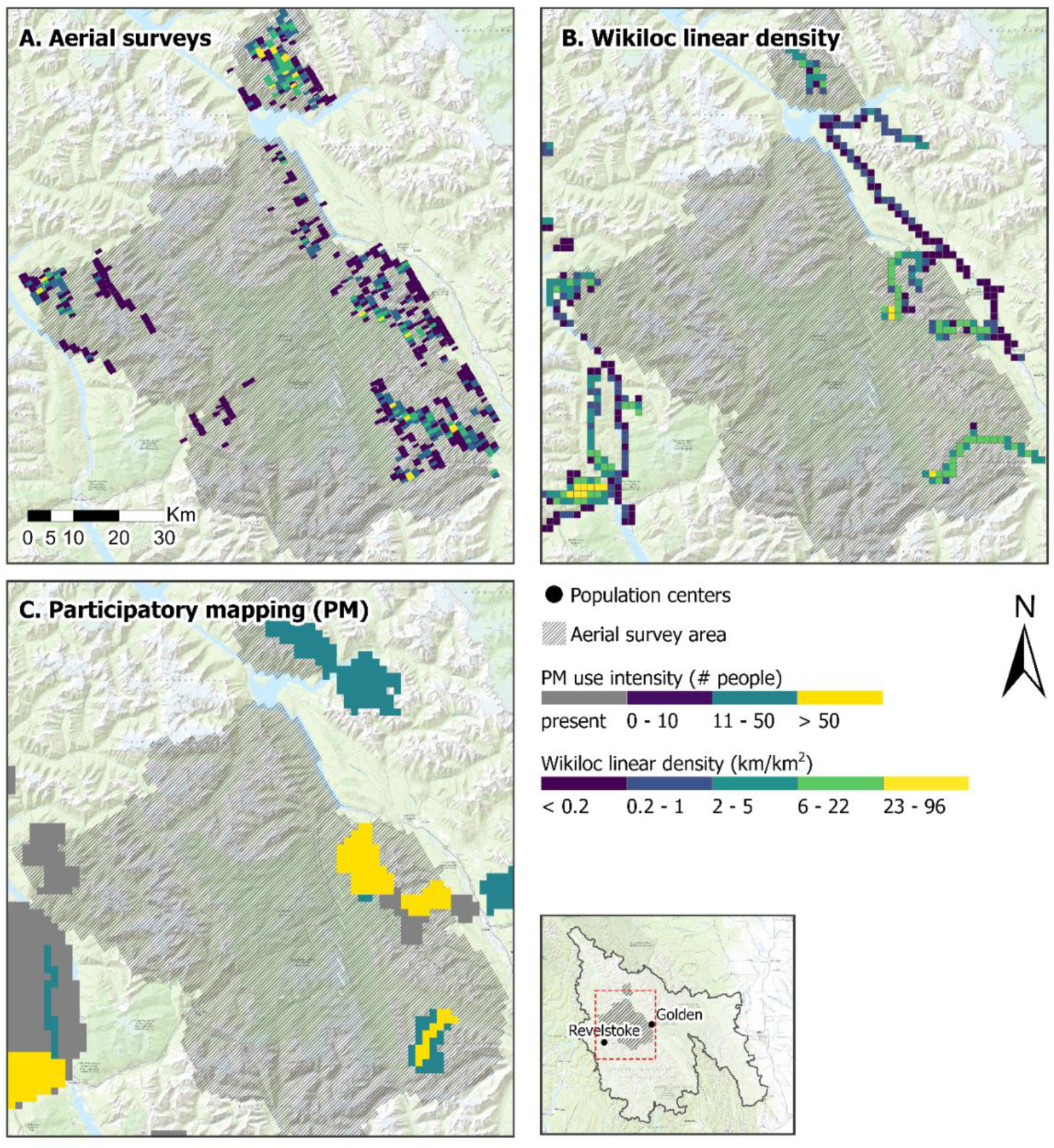
Winter motorized recreation intensity for aerial surveys (A), Wikiloc linear track density (km/km^2^; B), and participatory mapping (PM; C) in the Kootenay Mountains, British Columbia, Canada. Aerial surveys represent the percent footprint of snowmobile tracks in each grid cell. Hashed gray shading represents the areas surveyed by aerial flights between February and April 2022. Wikiloc linear density represents the total density (km/km^2^) of winter (November – April) GPS tracks from the app-users, 2015 – 2023. Participatory mapping represents the estimated number of motorized winter recreationists, 2017 – 2019.

All tools provided similar estimates of where winter motorized recreation occurred across the landscape, although patterns in the spatial extent and intensity of recreation use varied due to the resolution and activity type of the datasets (Fig. 8). PM identified areas where recreation was occurring over the broadest extent, but did not provide continuous or as detailed spatiotemporal information on the number of recreationists as the other tools. Although at a coarser resolution than the other tools, PM captured similar patterns as Wikiloc for areas with high winter motorized use. Comparatively, aerial surveys captured areas with snowmobiling that the two other tools did not, particularly in areas with low intensity of use (Fig. 8). This incomplete representation of winter motorized recreation by Wikiloc and PM, particularly in areas with low intensity of use, may be attributed to the application-based data being dependent on the number and types of users of the app and PM participants not being as familiar with less popular areas for motorized winter recreation.

While we demonstrate how traditional and application-based tools can collect off-trail winter recreation data, the accuracy of estimates, type of recreation activity, and spatial and temporal resolutions varies across tools. Combing multiple tools allows for areas with a range of use intensity to be represented and for more robust estimates of winter motorized recreation.

Moreover, this case study underscores the need for further development of tools to accurately monitor dispersed, off-trail winter-recreation at scales to match growing winter recreation footprints. This information is critical to better understanding and managing the impacts of winter recreation on sensitive and snow-associated species (Heinemeyer et al., 2019).

### 4.3 Canmore Biking

The Bow Valley of Alberta is a hub for year-round outdoor and adventure tourism and is a highly sought-after location for people to live, work and recreate (Alberta Government, 2018). Both humans and wildlife concentrate their activities along valley bottoms and as a result these areas are disproportionately affected by human development that can lead to human-wildlife occurrences, displacement of wildlife from habitats and loss of important wildlife corridors (Alberta Government, 2018). Mirroring many mountain towns in North America, the town of Canmore has undergone growth in their permanent population, development footprint and recreation sector (Alberta Government, 2018; Carlson et al., 2022), including the increase of the creation and use of informal recreation trails (Whittington et al., 2022). While monitoring and research on recreation patterns and impacts to wildlife has been identified as a research priority in the Bow Valley (Carlson et al. 2022), this work is hampered by lack of current and detailed information on human recreation activities. While traditional tools, such as cameras and counters, have been used to monitor recreation in this area, application-based data from fitness tracking apps has been largely unexplored.

Here we illustrate how the application-based tool Strava Metro provides similar estimates of bikers as cameras and counters on trails networks surrounding the town of Canmore. The three datasets were acquired from multiple research projects (see supplemental information for details) and were not always aligned across space or time. We therefore selected cameras and counters if were within 30 m of a Strava Metro trail segment and had data for the same day and referred to these as spatially matched locations. We calculated Pearson’s correlation values for monthly counts of bikers between Strava Metro and cameras or counters for all spatially matched locations from three years of data (2017, 2018, 2019). Correlations were plotted visually to identify spatial patterns in correlation (Fig. 9).

**Figure 9.**
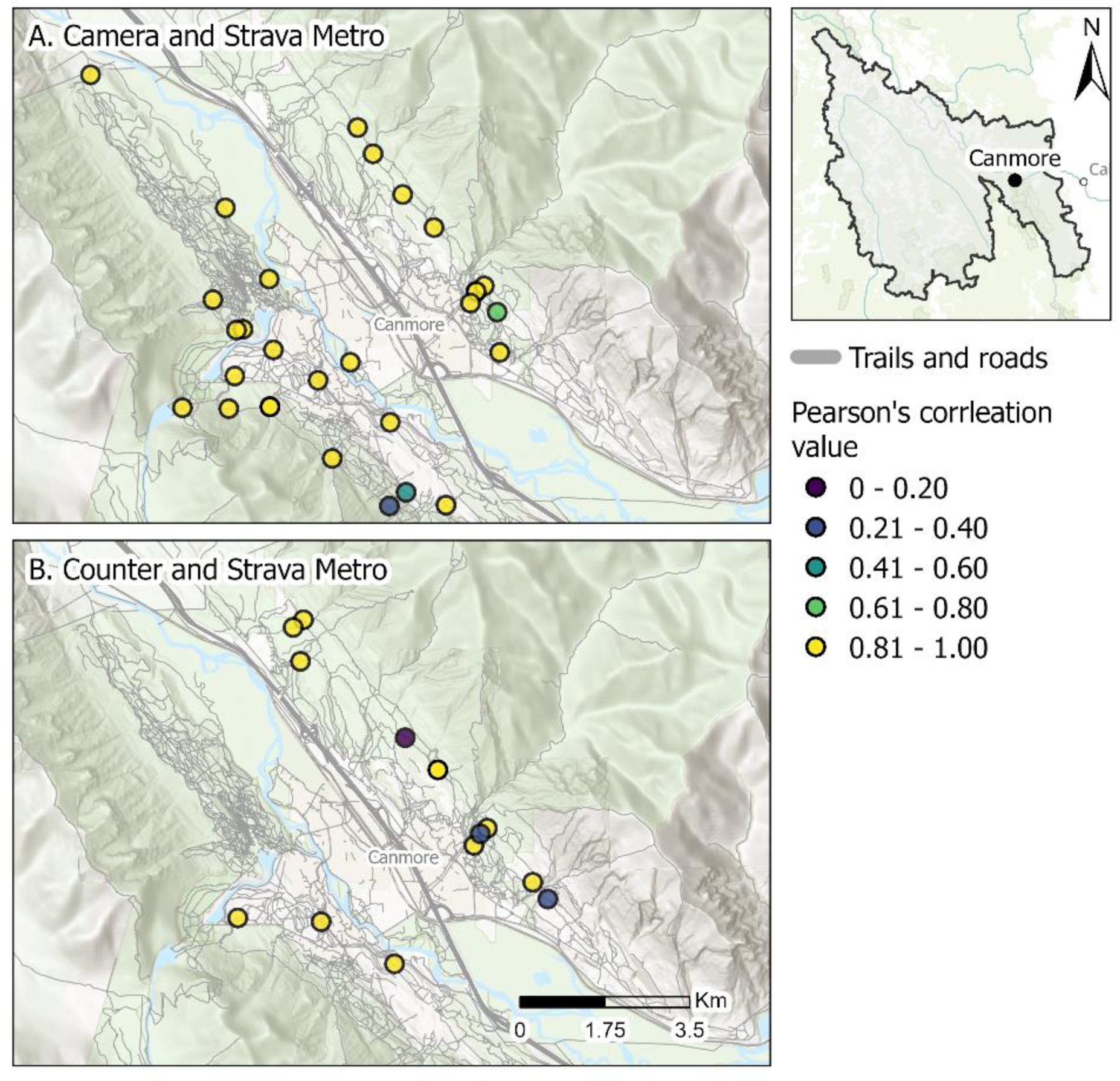
Pearson’s correlation values of monthly counts of biking activities for Strava Metro compared to cameras (A) or counters (B) in Canmore, Alberta, Canada.

Strava Metro captured similar monthly counts of bikers to cameras and counters (Fig. 9); 89% and 77% of the spatially matched locations of Strava Metro and camera or counter locations had high correlations (*r* ≥ 0.82), respectively (Fig. 9). These finding complements existing research demonstrating that Strava Metro data is generally a good approximation for the spatial extent of biking and use intensity (Lee & Sener, 2020). However correlation of Strava Metro to actual biker counts can vary depending on how data is aggregated spatially and temporally and influenced by user-group biases (Lee & Sener, 2020; Venter et al., 2023). Similar to many application-based recreation data, Strava Metro can overrepresent certain user groups (e.g., tech savvy, middle age, fitness focused; Venter et al., 2023). Canmore is a hub for recreation tourism and home to many recreation enthusiasts and therefore the high correlation among Strava Metro data to actual biker counts from cameras and counters may be attributed to a large proportion of bikers in the area using the app. However, as suggested by previous research (Venter et al., 2023), Strava Metro may not capture spatial and temporal patterns in biking across all local scales and consideration for the year-on-year increase in the platforms usership and types of users is also important for longer term analyses in patterns (i.e., over several years). Therefore, pairing Strava Metro with other tools, such as counters and cameras, allows for cross-validation and more accurate recreation estimates. For instance, the intensity of biking recreation could be estimated on trails without cameras or counters using Strava Metro and multipliers from other cameras and counter locations to account for Strava Metro representing counts of app users, not the volume of all bikers on trails. This approach allows for detailed recreation data to be scaled up to match growing recreation footprints and to meet the increasing demand for recreation data to guiding land use planning, species management, and sustainable recreation management.

## 5. DISCUSSION

The growth of recreation has come with a demand for tools that provide accurate, timely and detailed information on recreation use at scales that match expanding recreation footprints (Cessford & Muhar, 2003; Loosen et al., 2023). To make evidence-based decisions, managers and researchers require an understanding of the differences between tools and data sources. Our study supports the growing body of research illustrating how the combination of traditional and application-based tools expand the rigor and scope of estimating recreation use across large landscapes, although the accuracy and utility of each tool is context-dependent and varies across space, time and activity type. Cameras and counters captured similar broad-scale patterns in count estimates of all recreation activities and pedestrians across the study area. Although application-based data provided detailed spatiotemporal information on recreation use, datasets do not represent the full recreation population and were biased towards specific recreation activities. For instance, Strava Metro was more suited for capturing broad-scale spatial patterns in biking than pedestrian recreation. Traditional tools including aerial surveys and PM captured recreation footprint and intensity of use. Aerial surveys were able to capture areas with low intensity of recreation and PM was more suited for gathering recreation information across large spatial and temporal scales. To address the need for guidelines outlining available recreation monitor tools, we offer a comparison of each tool, highlight their strengths and weakness and provide suggestions on how to use these tools to address applied monitoring and management scenarios (see Table S2 for a comparison across the tools).

Correlations between cameras, counters and Strava Metro varied across space, time, and activity type, highlighting the ability of each tool to capture specific activity types and users (Fig. 3; Fig 4). Compared to cameras and counters that captured activities regardless of if a recreationists was using an app, data from application-based tools reflect particular activities and a proportion of the total recreation population (Norman & Pickering, 2017). Previous research indicates that Strava Metro over-represents bikers relative to pedestrians (Venter et al., 2023), which may explain why we found that Strava Metro had a stronger correlation to cameras and counters for biking than for pedestrian activities across space (Fig. 3). In contrast, we found low correlation values for comparisons of cameras and counters for biking (Fig 3; Fig 4). This pattern is likely attributed to the low number of spatially matched camera and counter locations (Fig. 3) or counters underestimating or overestimating use. Counters cannot distinguish people in a group and may underestimate groups of recreationists, or overestimate use when counters are triggered by non-target movements such as moving vegetation and animals (Miller, Leung & Kays, 2017; Lawson 2021; McCahon, Brinkman & Klimstra, 2023). Pairing cameras, that provide spatiotemporal count information on group size and activity types, with counters in the same location would allow for counters to be cross-validated and their accuracy assessed.

We found clusters of lower correlation values for comparisons of Strava Metro to cameras and counters surrounding some population centers (Fig. S7) and during some summer months (May – August; Fig. 4) when activity counts peaked (Fig. S6). Lower correlations near populations centers were generally more pronounced for counter and Strava Metro companions for all recreation activities and pedestrians than bikers. This finding contrasts previous research demonstrating high correlations of Strava Metro and counters across space (Venter et al., 2023). A possible explanation for the trend we observed may be that the Strava Metro data does not represent the total recreation population or volume of recreationists in the study area, particularly near population centers and during the summer. The population centers in our study area are surrounded by high-density trail networks and are hubs for recreation tourism. Strava Metro is geared towards competition and fitness-focused users and displays spatial and user-group biases, with previous research finding larger proportion of the app users are in remote and low intensity use areas (Venter et al. 2023; see Lee & Sener, 2020 for review of Strava Metro). Further, Strava Metro does not represent children under the age of 13 (the minimum age to use the app; Strava Terms of Service 2023). Taken together, the lower correlations near population centers and in the summer may be attributed to differences in the behavior between fitness-focused Strava users who may travel further into remote recreation areas (Venter et al., 2023), and summer tourists and families that recreate closer to population centers and are less represented in the apps data.

We found generally lower correlation between biking counts for Strava Metro and cameras or counters with increasing temporal aggregation. Correlations were higher for the spatial comparison when monthly biking counts were aggregated across all years for each unique camera or counter spatially matched location (mean correlation values, *r* = 0.81, *r* = 0.82 for Strava Metro comparison to camera and counters, respectively; Fig. 3) compared to the temporal comparison when monthly biking counts were aggregated by year and there was more variable correlations (Fig. 4). These findings are similar to previous research demonstrating higher correlations between Strava Metro and counter stations across space than time (Venter et al. 2023). As suggested (Venter et al. 2023), this trend may be attributed to Strava Metro not including all data at higher temporal units of aggregation (i.e., daily) because the privacy threshold requires at least three activities per unit time to be stored on a database. As a result, monthly or annual aggregation may allow greater number of trail segment to be included, particularly in areas less popular for recreation by Strava users.

Strava Heatmap is a popular index of cumulative annual recreation intensity and is increasingly used as a proxy for human recreation (Jäger, Schirpke & Tappeiner, 2020; Corrandi et al,. 2021; Carlson et al., 2022). Previous research demonstrates that Strava Heatmap reflects the intensity of ski mountaineer activities from counting stations (Jäger et al., 2020). In contrast, we found that Strava Heatmap did not consistently capture similar patterns as other tools: high recreation counts from cameras, counters and Strava Metro corresponded with both high and low Strava Heatmap index values, and cameras and Strava Metro captured recreation in locations where Strava Heatmap indicated no recreation. Strava Heatmap only reflects a proportion of recreation users and many not pick up all recreation users not using the app in less popular areas. We stress the importance of accounting for this limitation, especially when analyzing interactions between recreationists in less popular areas where there are sensitive, wary wildlife species, such as wolverines and caribou, whose habitats are often in remote and rugged terrain and who can be sensitive to very low levels of human-use (Barrueto et al., 2022). The mismatch across tools may also be attributed to temporal differences among our datasets. The Strava Heatmap dataset represented cumulative recreation data from 2016 to 2017, whereas cameras, counter and Strava Metro data represented recreation use across three years (2017, 2018, 2019). Furthermore, Strava Heatmap is not a straightforward index: map index values are not comparable at large distances because the same colour only represents levels of recreation intensity locally (Robb, 2017). At our Heatmap zoom level of 12 (190 m resolution), heat map values are comparable within an approximate 50 km diameter, similar to previous studies (Corradini et al., 2021). Our results suggest that the Strava Heatmap may be more appropriate to use for smaller extents, such as within our case study areas.

Our findings highlight how application-based products provide an incomplete representation of recreation because they only capture the relative popularity of recreation in areas among app users. For instance, in the Ghost Public Land Use Zone (PLUZ) case study, we found that application-based Wikiloc data did not capture all summer motorized trails. Moreover, application-based data exhibits unequal contribution of data among users because the most active users contribute more than others and is often biased towards demographic groups (Ghermandi & Sinclair, 2019, Venter et al., 2023). These comparisons highlight how the spatial and temporal distribution of application-based data are dependent on the number and types of users on the platform. This may be further compounded by changing recreational use patterns and app usership over time (Venter et al., 2023). While application-based tools can provide detailed data about recreation for large areas, the data may not reliably address spatial or temporal recreation patterns if biases in app use are not properly accounted for. As we demonstrate here, cross-referencing application-based data with traditional tools such as cameras or counters offers a solution for identifying these biases and improving the accuracy of recreation estimates.

In contrast to the application-based data from Strava products and Wikiloc, PM and aerial surveys provided information on where recreation was occurring regardless of activity or app use (Fig. 6). Our PM data was collected at a coarse spatial scale, making it challenging to directly compare with spatiotemporal data from cameras, counters, and GPS-track data from Strava Metro or Wikiloc across the study region. Similarly, the aerial survey data was difficult to compare with data from other tools because it represented a single snapshot of recreation use in time. Although multiple surveys can be conducted throughout the season to increase the temporal coverage of aerial survey data, the cost and time for aerial surveys limits how frequently this type of data can be collected. An advantage of Wikiloc, PM and aerial surveys is that these tools collect off-trail recreation data without having prior knowledge of where recreation occurs.

Aerial surveys capture areas with low intensity of use that other tools did not (Fig. 8) and when combined with other tools can provide more robust and detailed recreation information.

### 5.1 Study Limitations

While our work provides insight into the application of traditional and application-based tools for monitoring recreation, our results need to be interpreted in light of the study’s limitations and scope. Our datasets were collected from various wildlife and recreation monitoring projects, resulting in a spatial and temporal mismatch between datasets. Further studies would benefit from having tools with the same spatial and temporal coverage to allow for more direct comparisons of tools across greater areas and time periods. Despite this, our approach of leveraging multiple datasets from existing projects reflects data collection limitations often faced by managers due to time, funding, and staffing constraints. Further, this approach demonstrates how supplementing data from other projects can alleviate the costs and labor associated with data collection and can enhance the temporal and spatial extents and sample size of data.

### 5.2 Future Research and Opportunities

Our study highlights where more research is needed. While our work generally focuses on recreation on trails, recreationists continue to venture off-trail and further into the backcountry. Our research on aerial surveys and Wikiloc adds to previous research capturing off-trail recreation (Norman & Pickering, 2017; Heinemeyer et al. 2019). Moreover, due to the datasets available, we only assessed the ability of Wikiloc to capture motorized recreation in specific locations. Because Wikiloc reflects recreationists in more remote areas (Norman & Pickering, 2017) and captures a variety of recreation activities, future work should assess the platform’s ability to quantify off-trail use for various activities to better understand how to maximize the benefits of this tool. However, use of this tool should incorporate careful consideration of biases in the datasets and constraints of manually downloading each individual Wikiloc track (Goodbody et al., 2021). Alternatively, traditional PM holds promise for identifying broad areas with informal recreation trails and activities not readily captured by other tools, such as winter and water-based recreation. Information from PM could be used to strategically select areas for ground or aerial surveys, camera or counter placement, or prioritize areas for collecting application-based data.

By leveraging multiple datasets from existing projects, we addressed the call for more collaborative research and data sharing (Parr & Cummings, 2005; Steenweg et al., 2017). Numerous networks have been formed to facilitate the use of monitoring tools and data collaboration in ecology research (e.g. DataBasin; WildCAM; WildTrax; Steenweg et al., 2017). Development of a specific human recreation community of practice (e.g., https://wildcams.ca) would help standardize and centralize recreation data and promote data sharing to improve monitoring and management. Further, collaboration and standardization of monitoring and data practices would help address the challenges of insufficient staff and financial resources limiting the adoption of application-based data for recreation monitoring and management (Mangold et al., 2023).

### 5.3 Implications and Recommendations for Management

Identifying tools that provide accurate and detailed data on recreation use is the foundation of recreation monitoring and sustainable recreation (Cessford & Muhar, 2003; D’Antonio et al., 2012). Selection of the use of a single, or combination of tools, should be based on an understanding of the strengths and limitations of each tool relative to clearly defined management and planning objectives and priorities (Cessford & Muhar, 2003). We provide a summary table (Table S2) and highlight the key strengths and limitations of the seven tools we assessed.

Evidence-based decisions often require detailed and context-specific information about recreation use and spatial patterns. For managers seeking to quantify fine spatial and temporal patterns of on-trail recreation within a park or trail network, cameras and counters can provide accurate and detailed information. The decisions to use camera traps or counters should be based on context-specific considerations. For instance, if managers require information on the type of recreation activity and group sizes, cameras should be used rather than counters because the latter lacks the capacity to differentiate across all recreation types and group sizes (Table S2).

While cameras provide detailed data on recreation types and number of recreationists, this tool requires consideration of privacy and is resource intensive (e.g., replacing batteries, processing images, but see Fennell, Beirne & Burton (2022) for details on automatic classification software to improve image processing efficiency). It is also worth noting that cameras can monitor wildlife and population trends, potentially facilitating research on human-wildlife interactions to guide land-use management and planning (Steenweg et al., 2016; Naidoo & Burton, 2020; Barrueto et al., 2022). Comparatively, data processing for counters is generally quick and has low labor requirements (Miller, Leung & Kays, 2017), making this tool practical for long-term monitoring. However, counters systematically underestimate recreation use when groups of people trigger the counter only once (Miller, Leung & Kays, 2017; Lawson, 2021) and can overestimate recreation use when triggered by animals or vegetation. We suggest that when possible, pairing tools, such as counters with cameras, can derive additional data, allowing for cross-validation and more accurate recreation estimates.

Application-based tools, such as Strava Metro and Wikiloc, can supplement traditional tools and provide detailed information across large landscapes, although these platforms vary on the types of human activities reflected in their data. Use of application-based tools must be carefully considered, with biases in spatial extents and user groups addressed (Table S2). Thus, we suggest that application-based data be supplemented with traditional tools such as cameras or counters or data from regional or large-scale monitoring efforts already underway to improve confidence in recreation estimates. Alternatively, when detailed recreation data at fine spatial resolutions is not needed, but extent is important, aerial surveys, PM and Strava Heatmap offer alternative approaches to collect coarser recreation data. Strava Heatmap expands the scope of recreation activities and includes winter and water activities in addition to hiking, walking, running and biking. However, the information from this platform is at a coarse time scale with maps representing annual cumulative index of recreation intensity, making it challenging to integrate with higher resolution data from cameras or counters. Similarly, aerial surveys can capture winter off-trail recreation, both extent and index of use. These surveys can be used to monitor changes over time in a repeatable methodology. However, weather, cost and labor may limit aerial surveys spatial and temporal extents. Alternatively, PM can be designed to provide information on intensity of use and recreation activity type with the desired spatial or temporal resolution to meet practitioners or manager’s needs. However, the quality of PM data is sensitive to sampling design, participants cognitive bias and their characteristics, knowledge of place, and experiences that shape perspectives (Brown, 2011; Brown & Kyttä, 2014). These attributes may limit the accuracy of PM data and need to be properly accounted for. Collectively, PM and Strava Heatmap are helpful in instances when there is little information on recreation use in an area. These tools can provide information on where recreation is occurring that can subsequently be used for prioritizing areas for placement of traditional tools - such as cameras and counters - to collect finer scale recreation data. The use of these three tools may be particularly helpful for cases when employing large networks of cameras or counters is impractical, such as monitoring off-trail recreation in large and rugged landscapes or across multiple parks or recreation areas. While incorporation of application-based tools for visitor monitoring and management is likely to increase in the future (Mangold et al., 2023), digitalization of outdoor recreation and visitor management presents challenges and opportunities for managers of recreation areas. Many recreationists use digital outdoor and fitness platforms for planning and navigating recreation activities. These tools can be used for visitor management (i.e., promoting official trails, adding trails information on crowdsourced map-services such as OpenStreetMaps; Mangold et al., 2023). However, unofficial trails (i.e., trails not managed for recreation) are frequently included in these platforms and are subsequently shared with many users (Mangold et al. 2023; Schwietering et al., 2023). These unofficial trails are often not included in government databases, particularly those from OpenStreetMaps that is the basis for many smartphone applications (Loosen et al., 2023). The lack of information about which recreation activities are occurring on unofficial trails is challenging for managers; managing human-wildlife conflict and conflict between recreation users is difficult if the extent of recreation and types of recreation occurring is unknown (Loosen et al., 2023). Leveraging information from application-based tools that monitor unofficial and off-trail recreation can provide managers with additional information to supplement and validate their current databases to build a more comprehensive understanding of the recreation footprints that underlies many decision-making and planning processes.

## 6. CONCLUSIONS

The growth of outdoor recreation has come with a demand for tools that accurately estimate the extent and intensity of recreation use. This information constitutes the basis for understanding the impacts of recreation on wildlife and natural areas, developing evidence-based land use management plans and promoting enjoyable and sustainable recreation. Yet, there remains a lack of timely, accurate, and detailed recreation data at spatial scales that match expanding recreation footprints. We addressed this knowledge gap by comparing the use of traditional and application-based tools for measuring recreation across a large landscape and local trail networks and highlight the strengths and weakness of each tool. Our research illustrates that recreation use can be estimated using traditional and application-based tools, although their accuracy and utility varies across space, time and activity type. We recommend that the specific context and management objectives guide selection of tools. Application-based data from apps such as Strava products or Wikiloc should be supplemented with data from traditional tools such as cameras or trail counters, to identify individual tool biases and improve confidence in recreation estimates.

Our research contributes towards a better understanding of what tools are available and can help direct managers in selecting which tool, or combinations of tools, to use to improve recreation use estimation and monitoring. This information can guide decisions to better protect ecological systems while allowing for sustainable recreation.

## Supporting information

Supplemental data description, tables and figures

## ACKNOWLEDGMENTS

We thank all data partners (Ministry of Forestry and Parks, Nature Conservancy of Canada, Parks Canada, Recreation Sites and Trails BC) for sharing data for this research and for the insight and feedback that guided earlier drafts of this paper. Special thanks to Naia Noyes-West for leading the participatory mapping and Kim Heinemeyer and Peggy Holroyd for thoughtful feedback and engaging discussion that helped shape this research. We also thank Loosen Studio for creating visual representations of our research. Thank you to the journal editor and reviewers for their constructive comments.

## AUTHOR CONTRIBUTIONS

Conceptualization: A.L; A.L.J; B.A.M.; K.P; L.E; P.W; T.V.C. Project supervision: A.L.J; L.E; P.W. Methodology: A.L; B.A.M; K.P; T.V.C. Data analysis: B.A.M; T.V.C. Writing original draft: B.A.M; T.V.C. A.F contributed data and all authors provided substantial editing to the final draft.

## DATA AVAILABILITY

We will share R code and a subset of this data upon publication on a public repository (e.g. Dryad, GitHub). Some datasets may not be available, or spatial coordinates will be withheld, due to security and the sensitive nature of some of the datasets.

