## Supplemental data description, tables and figures for "Advancements in monitoring: a comparison of traditional and application-based tools for measuring outdoor recreation"

### Recreation data sources: data collection methods

#### *Camera trap*

We compiled camera trap data shared from Parks Canada and Alberta Parks remote camera trap networks (Steenweg et al., 2016; Stewart et al., 2016; Whittington, Low & Hunt 2019). Recreation included motorized (motorbike, quad, snowmobile, vehicle) and non-motorized (angler, bike, climb, electric bike, fat bike, hike, hunt, run, ski, skijorer, snowshoe) activities. We categorized activity types into three categories: all activities (all motorized and non-motorized activities), pedestrian (angler, climb, hike, hunt, run, ski, skijorer, snowshoe) and biking (bike, electric bike, fat bike). A summary of camera sampling design and image processing approaches for each project are detailed below.

*Steenweg R, Whittington J, Hebblewhite M, Forshner A, Johnston B, Petersen D, Shepherd B, Lukacs PM. 2016. Camera-based occupancy monitoring at large scales: power to detect trends in grizzly bears across the Canadian Rockies. Biological Conservation 201:192–200. DOI: 10.1016/j.biocon.2016.06.020.*

Steenweg et al. (2016) monitored changes in the occupancy of grizzly bears (*Ursus arctos*) using camera-based occupancy models across the Canadian Rockies. They deployed 183 cameras per a 10 x 10 km cell across the Canadian Rockies study area including Jasper, Banff, Yoho, Kootenay and Waterton Lakes National Parks in 2012. They used mainly covert motion-trigger cameras (Hyperfire and Rapidfire models; Reconyx, Holmen, Wisconsin) and some visible glow cameras (Silent-image, Reconyx). Sites were selected to where animals were likely to travel based on topography, confluence of wildlife trails and grizzly bear rub trees. Cameras were set at waist height, pointed slightly downwards, and set to take five images per movement trigger with no delay between triggers. No bait was used and cameras were serviced 3-4 times a year. The Timelapse2 software (Greenberg and Godin, 2015) was used to classify images into species, sex and age classes. Picture events (i.e., independent detection events) were identified as images of grizzly bears at least 5 minutes apart.

*Stewart FEC, Heim NA, Clevenger AP, Paczkowski J, Volpe JP, Fisher JT. 2016. Wolverine behavior varies spatially with anthropogenic footprint: implications for conservation and inferences about declines. Ecology and Evolution 6:1493–1503. DOI: 10.1002/ece3.1921.*

Stewart et al. (2016) used camera traps to examine spatial patterns in the behavior of wolverine (*Gulo gulo)* across three study areas in the Rocky Mountains of Alberta, Canada. They deployed 164 Reconyx digital cameras (models RM30, PM30, PC900; Reconyx, Holmen, Wisconsin) in a systematic design composed of 12 x 12 cell grids between December and April, 2017 – 2013. Sites were baited. Cameras were programmed to take 5 photographs at 1 second intervals, repeated at each detected movement. We received this camera trap data categorized in a table, displaying for each camera image, the species, the total number of individuals in an image, and the date and time.

*Whittington J, Low P, Hunt B. 2019. Temporal road closures improve habitat quality for wildlife. Scientific Reports 9:3772. DOI: 10.1038/s41598-019-40581-y.*

Whittington et al. (2019) assessed response of nine mammal species to temporal road closures in Banff National Park, Canada, using a combination of remote cameras, road surveys and movement data from grizzly bear (*Ursus arctos*) global position system collar data. Motion triggered cameras (Reconyc Hyperfire PC900 Professional cameras, Holmen, Wisconsin) were part of Parks Canada broader remote camera network. Data was selected from 10 cameras along the Bow Valley Parkway and 54 reference cameras (18 on trails and 36 on highway crossing structures) from 2014 – 2017. Independent detection events were classified as whether or not at least one wildlife species was detected for each camera sampling day and hour.

***Trail counters***

We compiled data from passive infrared TRAFx trail or vehicle counters (TRAFx Research Ltd., Canmore, Alberta, Canada) shared with us from the government of Alberta, Parks Canada, Nature Conservancy of Canada (NCC) and Recreation Sites and Trails BC (RSTBC). Trail counters were deployed to monitor human recreation activity. In Kananaskis Country, Alberta, Canada a selection of these trail counters were paired with camera traps to examine differences in counts between the two tools to determine if the counters were installed in suitable locations and collected accurate data (Alberta Parks, personal communications).

***Participatory mapping (PM)***

Participants were selected from a list of recreation experts (i.e. park rangers, recreation group and trail users, lodge and campground owners) known to Yellowstone to Yukon Conservation Initiative (Y2Y). Participants were recruited by email with an explanation of the Y2Y recreation ecology research project (Fig. S1) and the areas where the project was interested in understanding human recreation use patterns (Fig. S2).

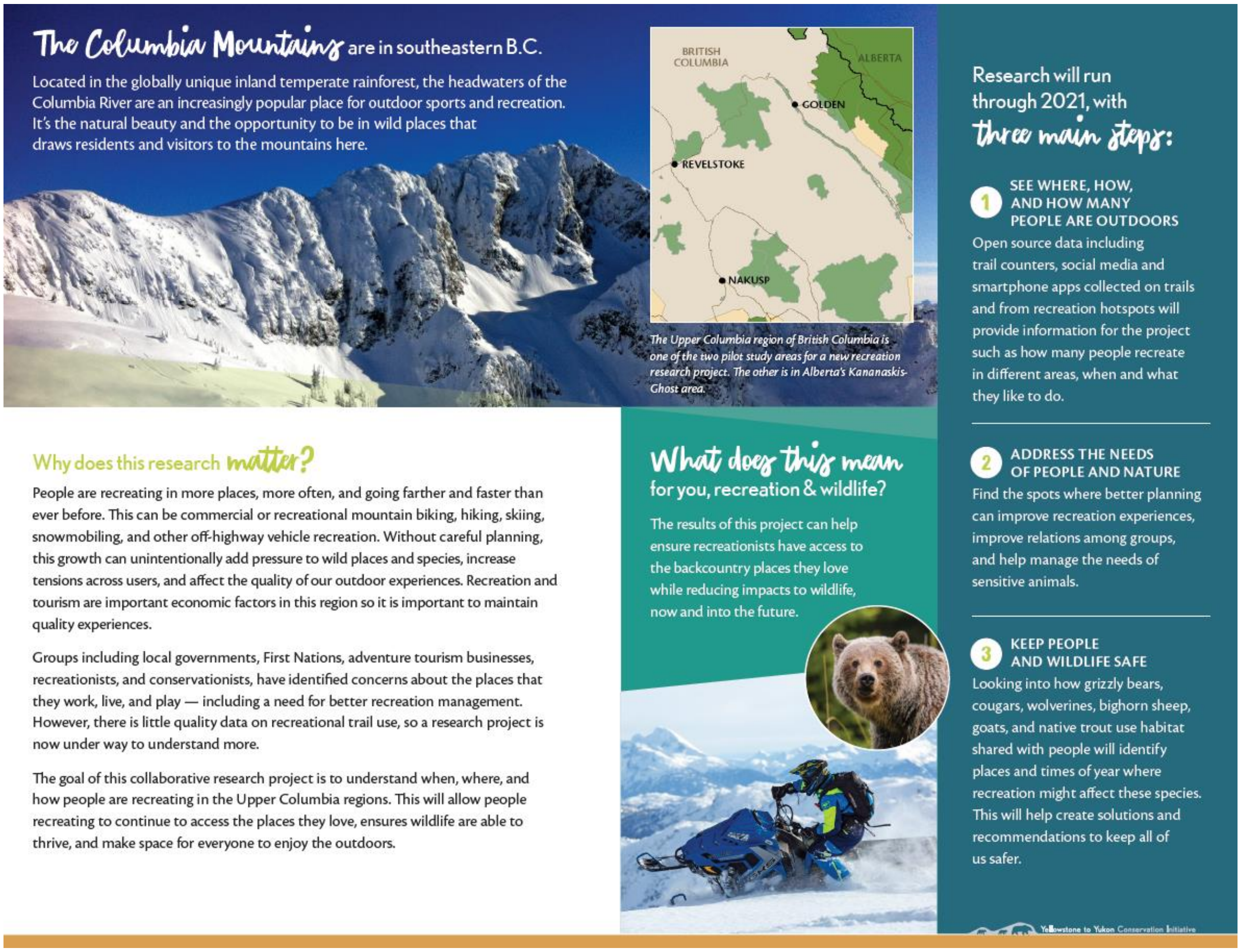


**Figure S1. Informational pamphlet sent via email to recruit volunteers in participatory mapping research to support the Yellowstone to Yukon Conservation Initiative recreation ecology project**. The image, and associated context, is out of date and reflects the early stages of the multi-year research project.

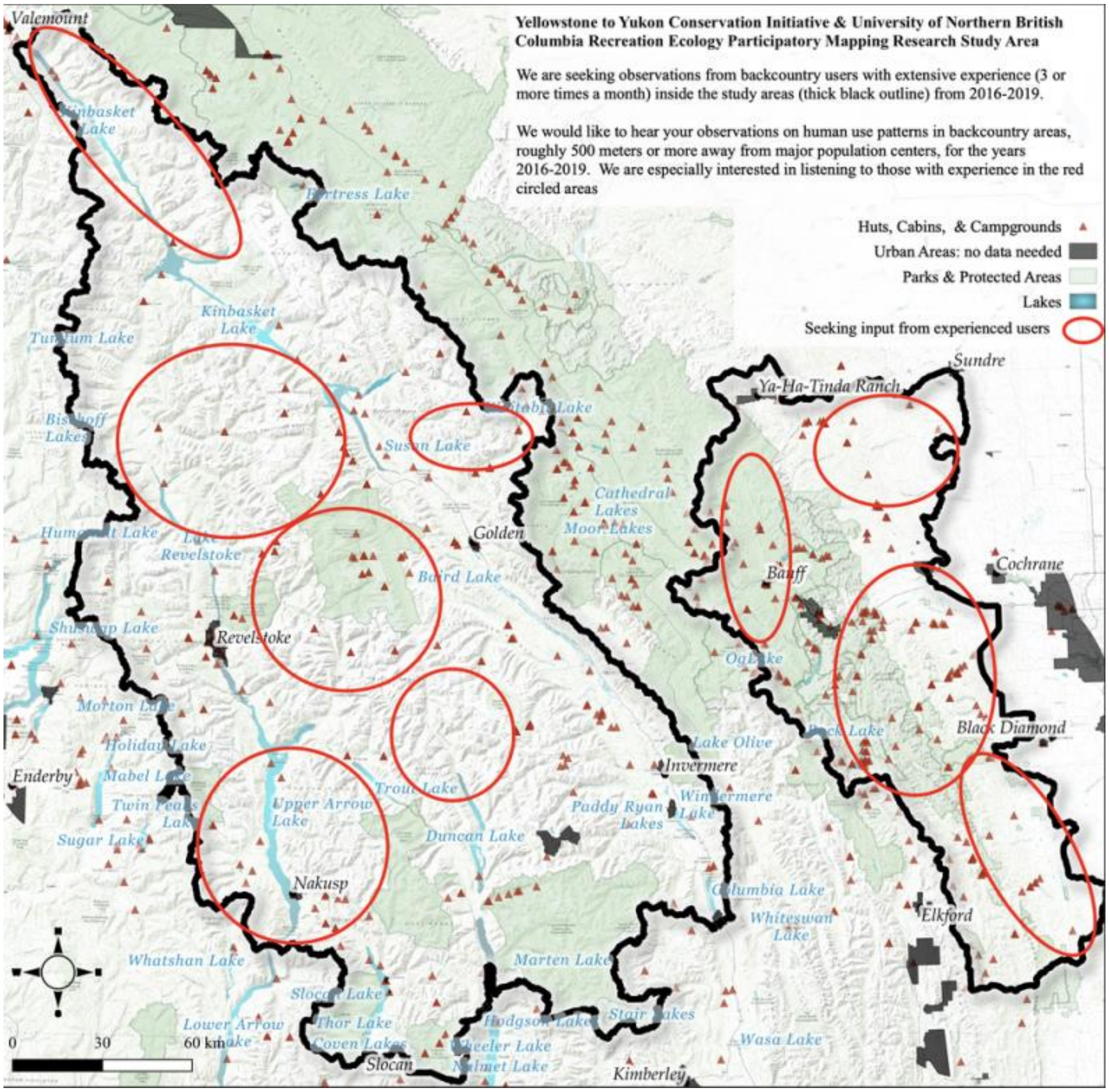


**Figure S2. Image of areas we were interested in collecting participatory mapping data sent via email to recruit volunteers in participatory mapping research to support the Yellowstone to Yukon Conservation Initiative recreation ecology project.** The image reflects the early stages of the multi-year research project. While the image indicates that we were interested in human use patterns for the years 2016 – 2017, we actually asked for information for the years 2017 – 2019. The study area in the image reflects the original study area boundary that was changed for later years of the project.

Meetings with 36 participants were conducted online over Zoom (Zoom Video Communications, Inc. 2020, San Jose, California United States) and were recorded with the permission of participants from 2020 – 2021. We started with a brief introductory explaining the research project and study area, outlined how we were going to proceed with the mapping, and asked if there were any questions or concerns. Participants were asked to load the web-based platform Gaia GPS (www.gaiagps.com) and to share their screen displaying an interactive map. They were asked to use the interactive map to show areas they were familiar with and explain what the average recreational use of that area was between 2017 – 2019. We asked if they could estimate daily use from 1–10 people, 11–50 and over 50 people a day and if they could specify what activity occurred in the area. If participants could only indicate if recreation was present or use within other years, we collected that information as well.

Recordings of meetings were watched and areas where participants described as having recreation use were digitized into ArcMap or QGIS points, lines or polygons with the activity type and use intensity added to the attribute table. Spatial resolution of the polygons where participants indicated there to be recreation use greatly varied; some participants provided estimates of the number of recreationists on a specific trail or camping area, while other participants provided recreationists estimates for a valley, land use zone or a large portion of our study area. Similarly, how participants provided estimates of recreationists varied. Some participants included information on the number of recreations over a weekend or throughout the week, sometime differentiate between seasons (i.e., summer or winter) or did not specify a time frame. Where appropriate, estimates were adjusted to reflect estimated number of recreationists per day.

In ArcGIS we classified all polygons and polylines of recreation use by activity type and season – summer (no snow on ground), winter (snow on ground), spring (snow only in alpine), fall (hunting season) – and a unique identification number was added to each individual polygon or polyline to identify the feature. A fishnet of grid cells was created to overlap each activity type and the unique identification number of each underlying polygon or polyline was extracted to each grid cell. Recreation count estimates were binned into three categories (0–10; 11–50; > 51 recreationists per day) and the maximum bin values was joined to the grid cells.

***Aerial surveys***

We conducted aerial surveys (Heinemeyer et al., 2019*)* in three focal areas in Alberta and British Columbia: i) the Kananaskis area south of Canmore, ii) the Upper Columbia between Golden and Revelstoke in British Columbia (BC), and iii) the Lake Louise section of Banff National Park, Yoho and Kootenay National Parks, Canada (Fig. S3). We selected flight areas in consultation with provincial and federal biologists, conservation officers and public safety specialists. A grid of 1.5 x 1.5 km cells overlaid the flight areas (Heinemeyer et al., 2019). Flight lines followed every other grid cell, resulting in transects spaced 3 km apart (Fig. S3). To survey, we flew the transect lines in a helicopter at an altitude of 305 m (1,000 feet) at a speed of 145 km/hr (90 miles/hour; Heinemeyer et al., 2019). In the rear of the helicopter, two primary observers sat on the left and right sides, each responsible for searching for snowmobiling, skiing (heli-skiing, backcountry skiing and cat-skiing) tracks. Observers recorded the recreation activity type, the percent of the observation window covered by recreation tracks (i.e., the footprint; none, 1–10 %, 10–25%, 26–50%, 51–75% and 76–100%), and track age (new – tracks were set after latest snowfall; old – tracks are snow covered). GPS waypoints were collected on handheld units every 20 seconds, during which observers scanned a 1.5 x 1.5 km area on each side of the helicopter. This defined the observation window. The survey leader ensured the helicopter followed each flight line and called out waypoint numbers to indicate to the observers when a new observation window had started or ended (Heinemeyer et al., 2019). Permit numbers were Alberta Parks (#22-024) and Parks Canada (#LL-2022-41098; RAP #KO22-001).

To match point observations to a viewing window, we created irregular grids extending out 1.5 km and forwards from the waypoint at which the observation window started to the next sequential waypoint. If there was a change in bearing (e.g. due to going off-course to avoid a peak, strong winds), the irregular grids were always perpendicular to the flight line (i.e., line between two waypoints; Fig. S4).

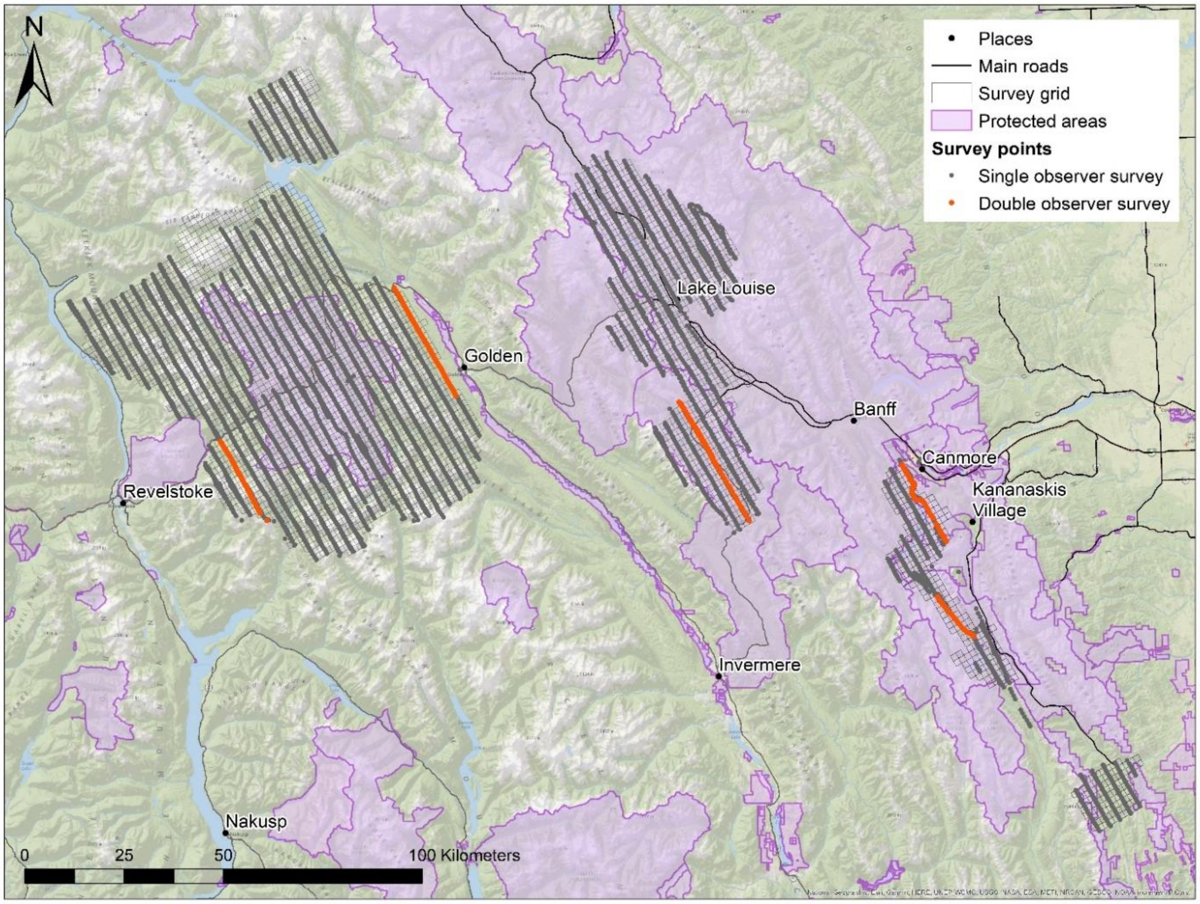


**Figure S3. Map of winter recreation aerial surveys in southeastern British Columbia and southwestern Alberta in winter 2022**. Grey dots represent single observer survey waypoints.

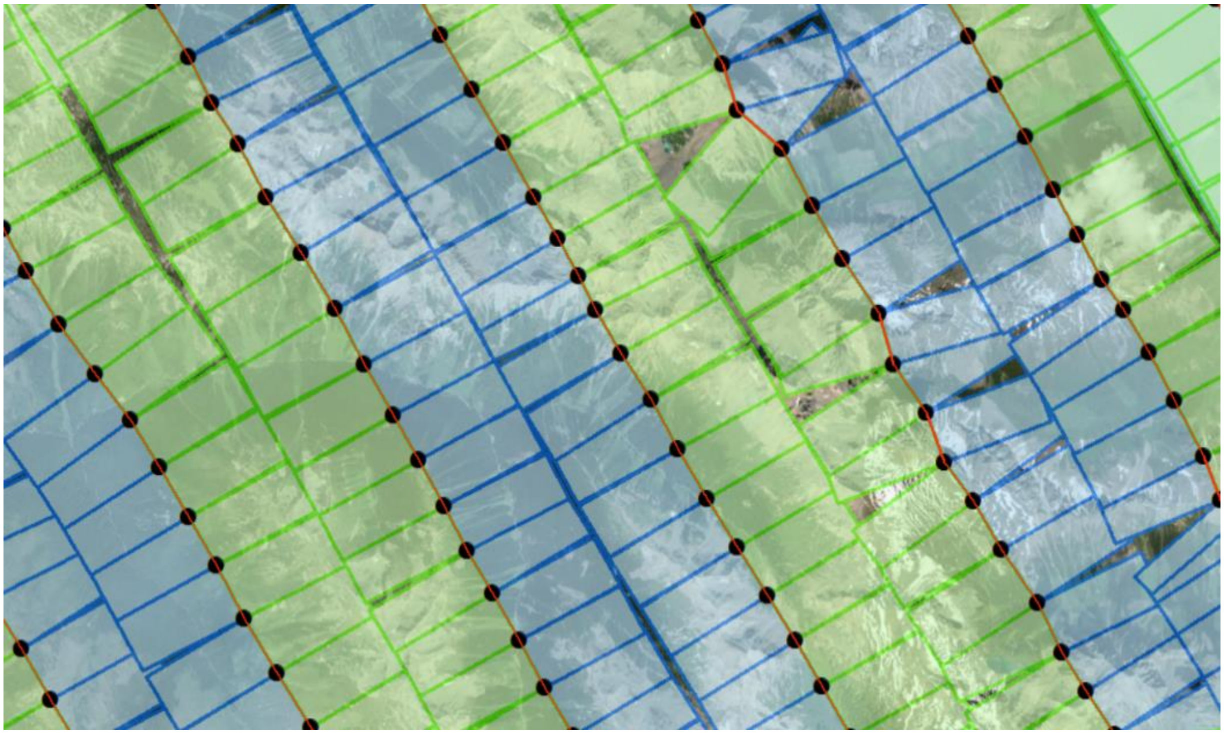

**Figure S4. Sample of an irregular grid for winter recreation aerial surveys.** Dots represent location of waypoints taken from the helicopter and the red line is the flight path of the helicopter. The blue boxes are the left-side observation window and green are right side observations. The grid length perpendicular to the red line is 1.5 km, representing how far out from the helicopter observers scanned for recreation tracks, while the width of the box was determined by the location of the following waypoint.

***Strava Heatmap***

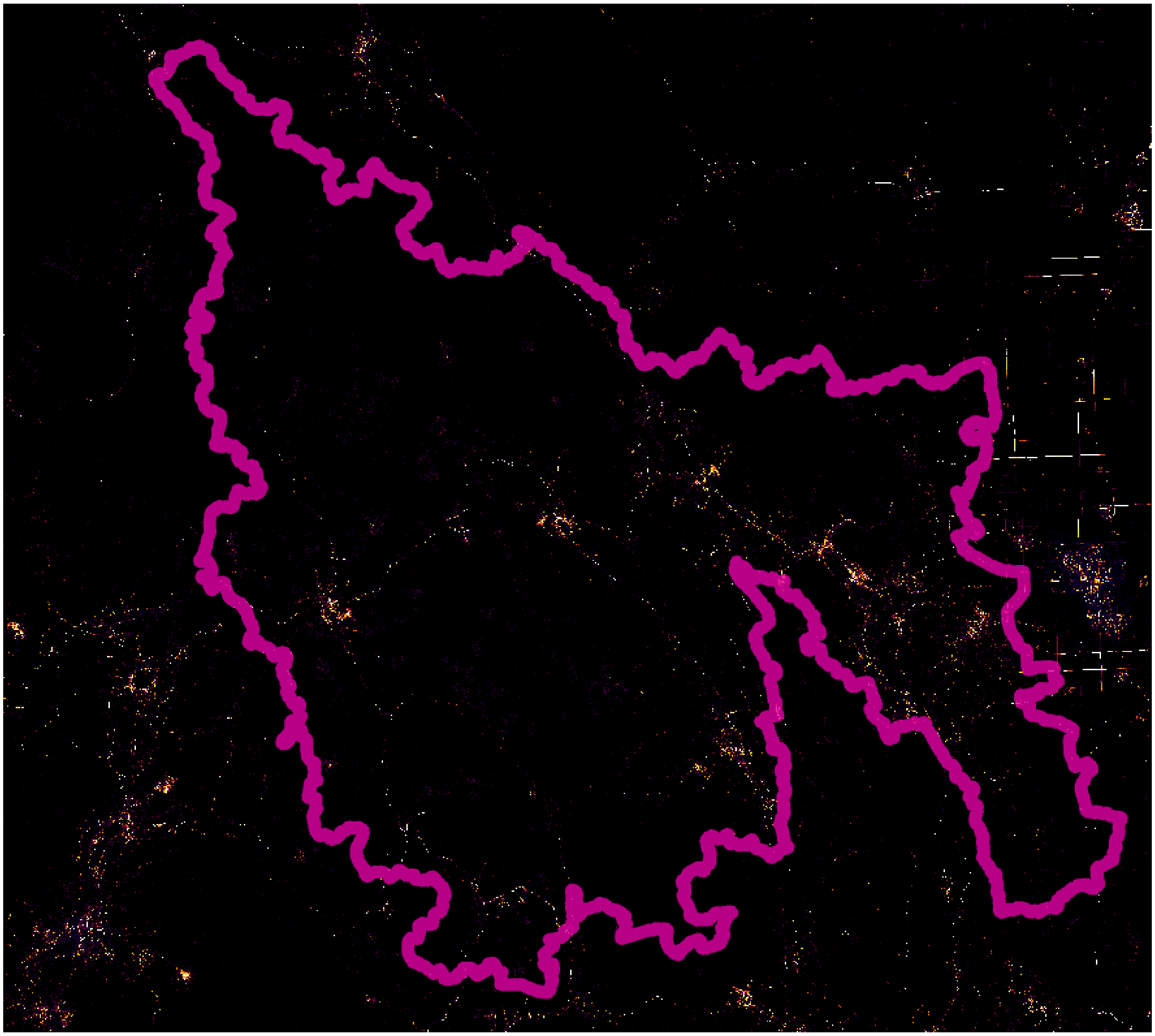

**Figure S5. Strava Heatmap from 2016 – 2017, with study area border denoted in pink.** Strava Heatmap represents cumulative biking, pedestrian, winter and water recreation activities from Strava users.

**Descriptive data summary**

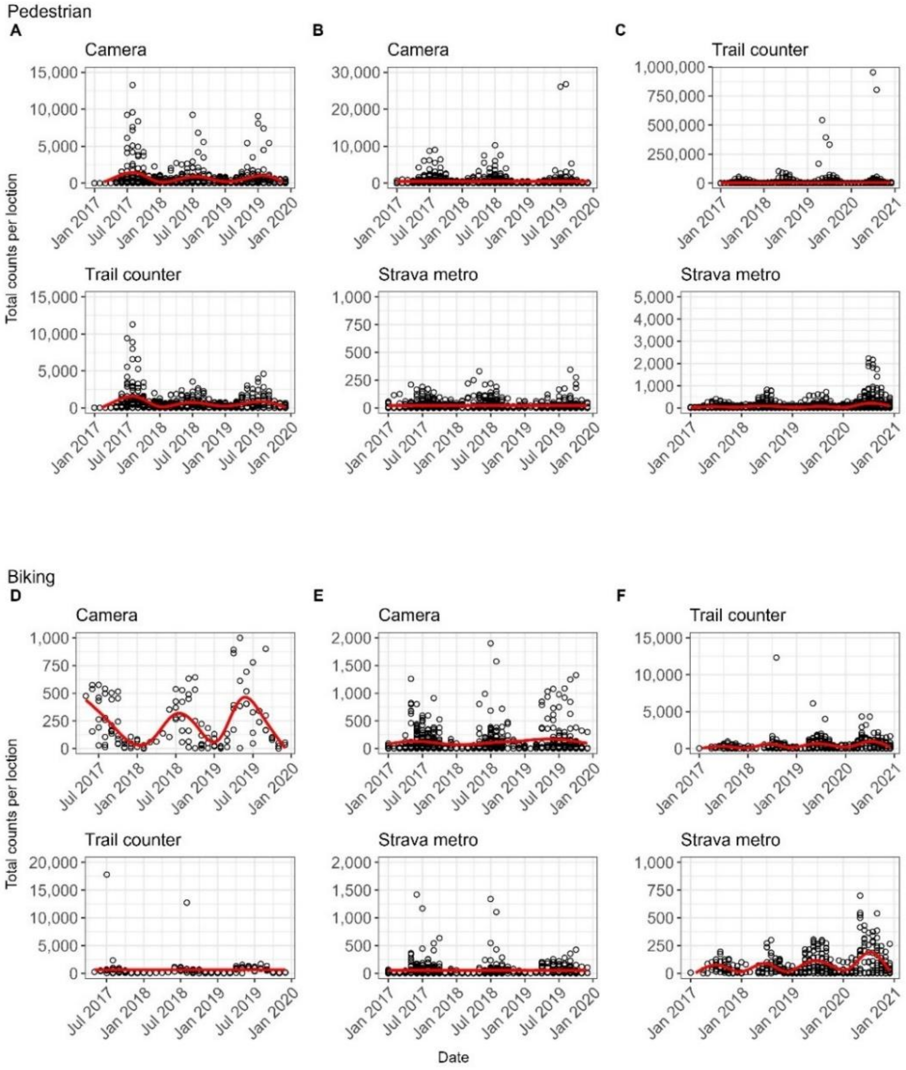


**Figure S6. Monthly counts of pedestrian and biking recreationists from spatially matched locations of cameras, counters and Strava Metro.** Each panel represents monthly pedestrian counts between spatially matched locations of (A) cameras and counters, (B) cameras and Strava metro, (C) counters and Strava Metro, and monthly biking counts from spatially matched locations of (D) cameras and counters, (E) cameras and Strava Metro and (F) counter and Strava Metro from 2017 – 2020. The red line represents a trendline generated with generalized additive models.

**Table S1. Number of pairwise combinations of camera, counter, and Strava Metro spatially matched locations for all recreation activities, biking, and pedestrian (hike, run, walk).** Cameras and counters were matched if devices were within 200 m of each other along a trail and had data on the same day. Cameras and counters were matched to Strava Metro segments within 30 m of devices and had data on the same day.

| **Activity type** | **Number of spatially matched locations** | | |
|---|---|---|---|
| | Camera and counter | Camera and Strava Metro | Counter and Strava Metro |
| All activities | 58 | 156 | 189 |
| Pedestrian | 60 | 157 | 184 |
| Biking | 11 | 102 | 36 |

**Comparison of monthly recreation counts across space for pairwise combinations of spatially matched cameras, counters and Strava Metro**

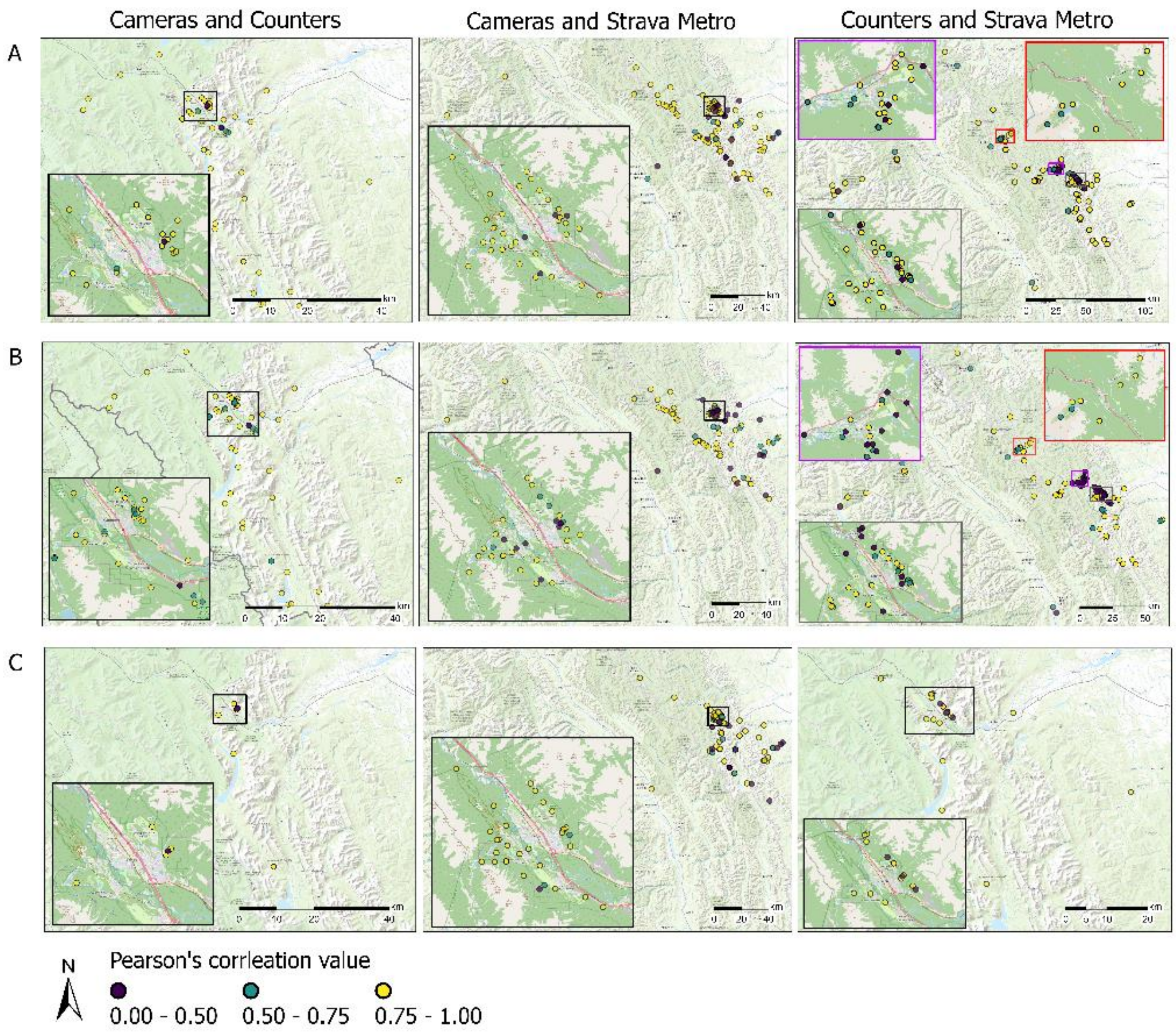


**Figure S7. Pearson's correlation values of monthly counts of all recreation activities (row A), pedestrians (row B), biking (row C) for pairwise combinations of spatially matched cameras, counters and Strava Metro locations.** First column compares counts from cameras and counters; second column compares counts from cameras and Strava Metro; third column compares counts from counters and Strava Metro.

**Strava Heatmap comparison to cameras, counters and Strava Metro**

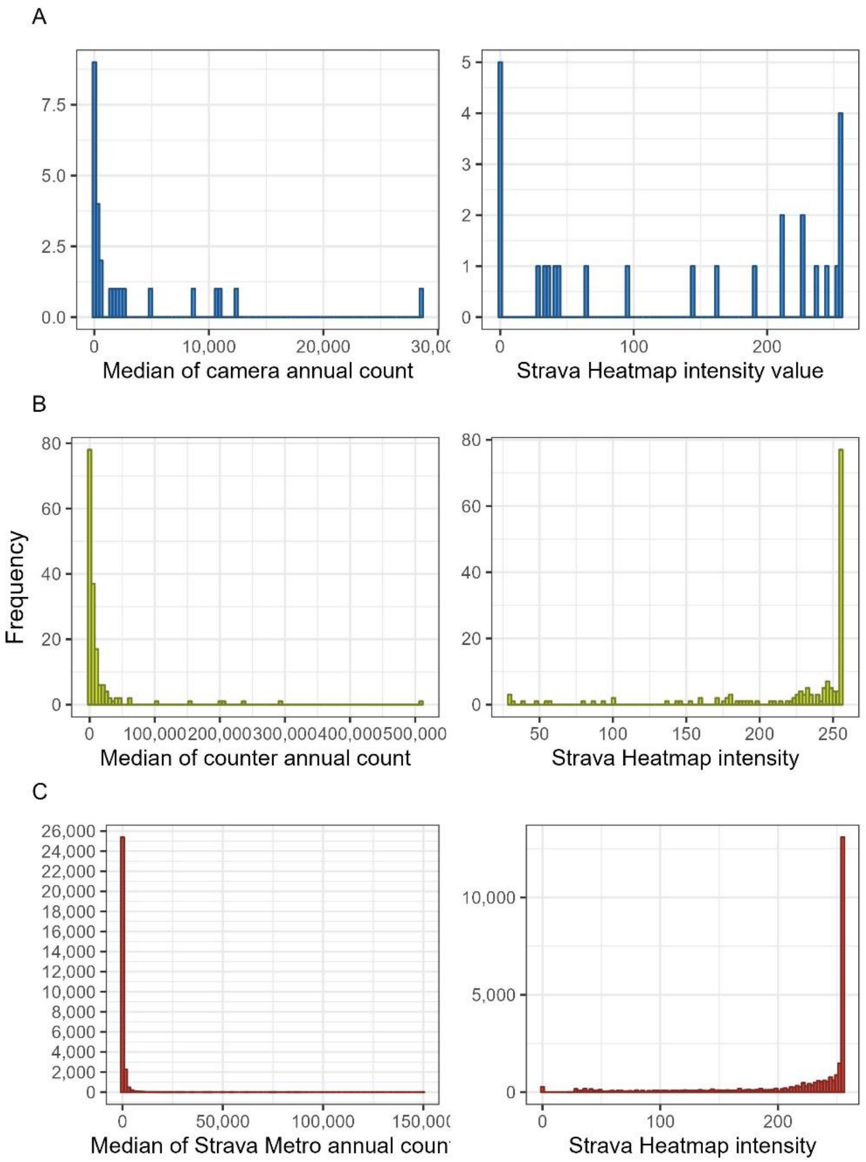


**Figure S8. Frequency distribution of the median annual counts of recreationists where cameras and Strava Heatmap values overlap (A; blue); counters and Strava Heatmap overlap (B; green); Strava Metro and Strava Heatmap overlap (C; red).**

**Spatial extent of motorized and non-motorized winter recreation from aerial surveys, Wikiloc and participatory mapping (PM).**

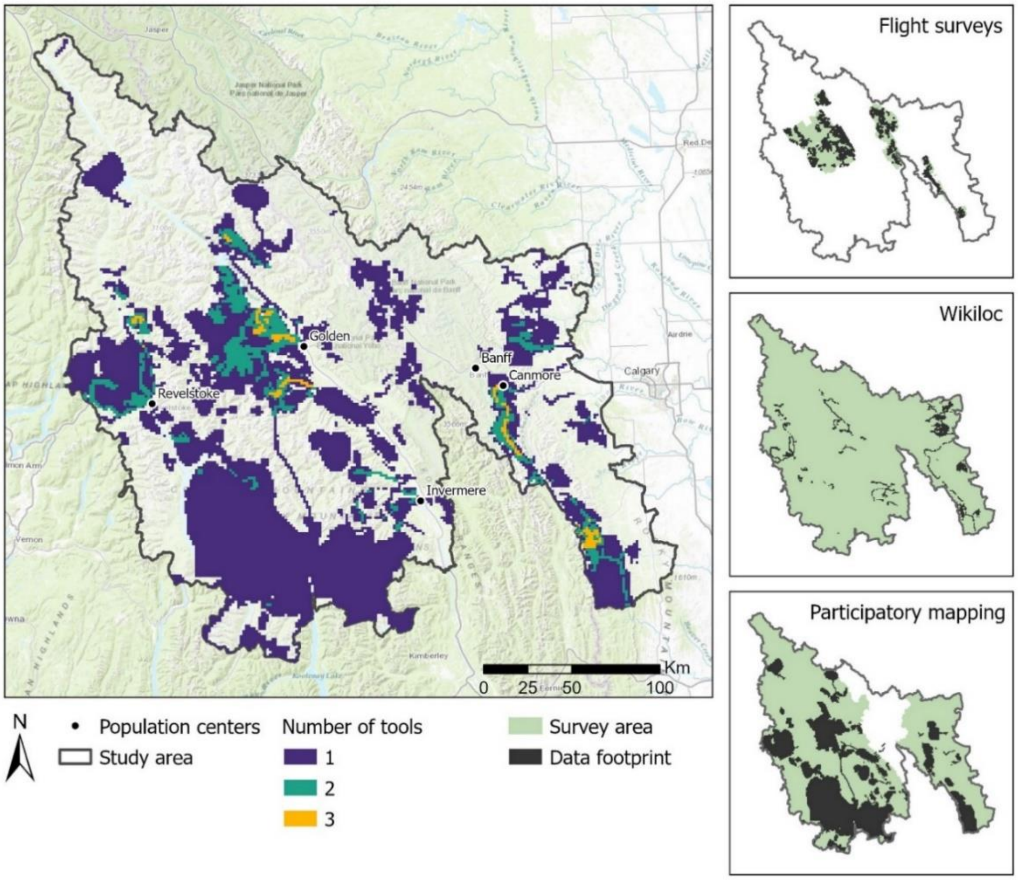


**Figure S9. Spatial extent of motorized and non-motorized winter recreation in the study area (left panel) from aerial surveys, Wikiloc and participatory mapping (PM).** Datasets from all tools were aggregated into 1.5 km cell grids. Panel A represents the number of tools where recreation was captured in the same 1.5 km cell. Only areas within the aerial survey were sampled for the aerial survey data (collected Jan – April 2022), a portion of the study area was sampled from the PM data (data representing recreation from 2017 – 2019) and the entire study area was sampled for the Wikiloc data (data representing recreation from 2015 – 2023).

**Summary of tools for measuring motorized and non-motorized recreation**

**Table S2. Comparison of the select monitoring tools to measure motorized (M) and non-motorized (NM) winter and summer outdoor recreation, including, but not limited to, key limitations and advantages for each tool.**

| Data or tool characteristic | Traditional tools | | | | Application-based tools | | |
|---|---|---|---|---|---|---|---|
| | **Trail counter** | **Camera trap** | **Aerial survey** | **Participatory mapping** | **Strava Metro** | **Strava Heatmap** | **Wikiloc** |
| **Counts of humans or GPS tracks** | Yes[1] | Yes | Yes | No | Yes[2] | No | Yes |
| **Spatial resolution** | High | High | Low to High[3] | Low to High[3] | High | High | High |
| **Temporal resolution** | High | High | Low to High | Low to High[3] | High[4] | Low[4,5] | High[4] |
| **Information on activity type** | Depends[6] | Yes | Yes[3] | Yes[3] | Yes[7] | Yes[8] | Yes |
| **Motorized (M) or non-motorized (NM)** | M and NM | M and NM | M and NM | M and NM | NM[7] | NM[8] | M and NM |
| **Off trail use** | No | No[9] | Yes | Yes | No[10] | Yes | Yes |
| **Data processing burden** | Low | Low to High[11] | Low | Low to High[3] | Low to High[3] | Low to High[3] | Low to High[3] |
| **Consideration for data privacy** | No | Yes | No | Yes[12] | Yes | Yes | Yes |
| **User-group bias** | No | No | No | No | Yes[13] | Yes[13] | Yes[13] |
| **Advantages** | | many references for setting up cameras, processing and analyzing data | continuous coverage of large areas; can capture photos and video; qualitative observations | qualitative information; can design methodology to capture areas of interests and level of detail | includes direction of travel | global coverage; includes more activities than Strava Metro (includes winter and water) | targets outdoor users in more remote locations; many recreation activities; can include photos and trip descriptions |
| **Limitations** | cannot differentiate between people, | difficult to capture off-trail use or trail | costly and weather dependent that can limit the spatial | sensitive to cognitive bias and misrepresentation of | biking and pedestrian | does not provide real counts of recreation; only | requires manually downloading each track |

| | | | | | | |
|---|---|---|---|---|---|---|
| | moving plants or animals, repeated counting if people stop in front of device; data error from environmental conditions and how people align themselves along trail; difficult to capture off-trail use or trail networks with numerous entry points; malfunction and tampering of device | networks with numerous entry points; malfunction and tampering of device | and temporal coverage; potential disturbance to recreationist and wildlife; potential observer biases; safety concerns | recreation on map; can be labor intensive to facilitate interviews and digitize paper maps or questionnaires; for large areas may have discontinuous coverage or incomplete information | recreation only | available at annual scale (not daily or monthly); heatmap values are comparable locally; can’t separate recreation activities (i.e., ski mountaineering, cross-country skiing, ice skating, etc.,); |

[1] Cannot differentiate between individuals or groups.

[2] Only includes activity counts (over 5 users) on trails in OpenStreet Maps.

[3] Can vary depending on context.

[4] Growing popularity with some recreation groups over time that need to be accounted for in temporal analyses.

[5] Only annual heatmaps are available (i.e., no daily or monthly heat maps are available).

[6] Some devices can specifically count bikes and cars.

[7] Includes only pedestrian (hike, walk, run) and biking activities.

[8] Aggregated into four recreation activity categories: water, winter, pedestrian (hike, walk run), and bike.

[9] Typically cameras placed to monitor recreation and wildlife trails or wildlife attractants (e.g., tree rubs for grizzly bears).

[10] All trails segments snapped to OpenStreetMaps street and trail segment.

[11] Advancements of automatic image processing software can speed up data processing.

[12] For participatory mapping participants.

[13] Unequal distribution of users, with some users contributing more data, more frequently
